# Xtrapol8: automatic elucidation of low-occupancy intermediate-states in crystallographic studies

**DOI:** 10.1101/2022.01.09.475568

**Authors:** Elke De Zitter, Nicolas Coquelle, Thomas RM Barends, Jacques-Philippe Colletier

## Abstract

Unstable states studied in kinetic, time-resolved and ligand-based crystallography are often characterized by a low occupancy, hindering structure determination by conventional methods. To automatically extract such structures, we developed Xtrapol8, a program which (i) applies various flavors of Bayesian-statistics weighting to generate the most informative Fourier difference maps; (ii) determines the occupancy of the intermediate states by use of *hitherto* unavailable methods; (iii) calculates various types of extrapolated structure factors while handling the issue of negative structure factor amplitudes, and (iv) refines the corresponding structures.

---

Once reserved to a handful of proteins and specialized laboratories, time-resolved and kinetic crystallography (TRX, KX) are on the verge of widespread adoption. This momentum is owed mostly to the advent of room-temperature serial crystallography, pioneered at X-ray free electron lasers (XFEL)^1^ yet swiftly implemented at synchrotrons where ease of access holds promise for groundbreaking studies.^2^ A main limitation of TRX and KX remains that full occupancy of the triggered state is hardly ever attained in the crystalline macromolecule, resulting in co-existence with the reference state. Low occupancy may also poison data collected from crystalline ligand-protein complexes, obscuring ligand identification and conformational changes undergone by the protein.

Based on the assumption that structure factor phases hardly vary upon reaction initiation and progression, Fourier difference electron density maps 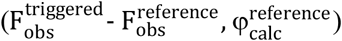 allow pointing to the largest structural changes between the reference and the triggered states.^3,4^ The information content of such maps can be improved by Bayesian-statistics (*q, k*) weighting of structure-factor amplitude (SFA) differences,^5,6^ yet they can be featureless when structural changes or the triggered-state occupancy are small, or if a mixture of triggered states^7^ – whose overlaying positive and negative difference peaks cancel each other – accumulate. Furthermore, Fourier difference maps allow modeling but not refinement of the triggered state structure. By use of extrapolation methods, whereby SFA differences are inversely scaled to the occupancy of the triggered state and summed with reference SFAs, the hypothetical SFAs for the triggered state present at full occupancy can be constructed, enabling both refinement of the triggered state structure, and to identify potential co-existing intermediate states 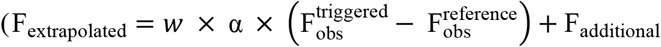; Methods and Supplementary Methods). With the advent of serial crystallography, the demand for extrapolation methods has risen yet these remain obscure to the vast majority of crystallographers for three main reasons. Firstly, the chosen occupancy is determined but hardly ever justified in absence of methods to estimate the triggered state occupancy based on crystallographic data only. Secondly, each lab resorting to structure factor extrapolation makes use of its own library of custom-written scripts, generally assembled over years of practice in TRX and KX, making results not easily reproducible. Thirdly, various flavors of extrapolation exist: exploiting or not Bayesian-statistics to downweigh SFA difference outliers; refining or not phases from the reference state structure against extrapolated SFAs (ESFAs); and recalculating or not the figure of merit of these phases prior to extrapolated map calculations. Here, we introduce Xtrapol8, a new software aimed at resolving these issues. Fig. 1 delineates its design and recommended usage. Below, we showcase its usage by application to recently published KX data collected on mEos4b, a photoconvertible fluorescent protein that emits green light in the ground state but can be converted to a red-emitting state upon UV illumination.^8^ In the Supplementary Information we further evaluate the versatility of Xtrapol8 by revisiting a variety of previously-published challenging TRX or KX studies.

**Fig. 1.**
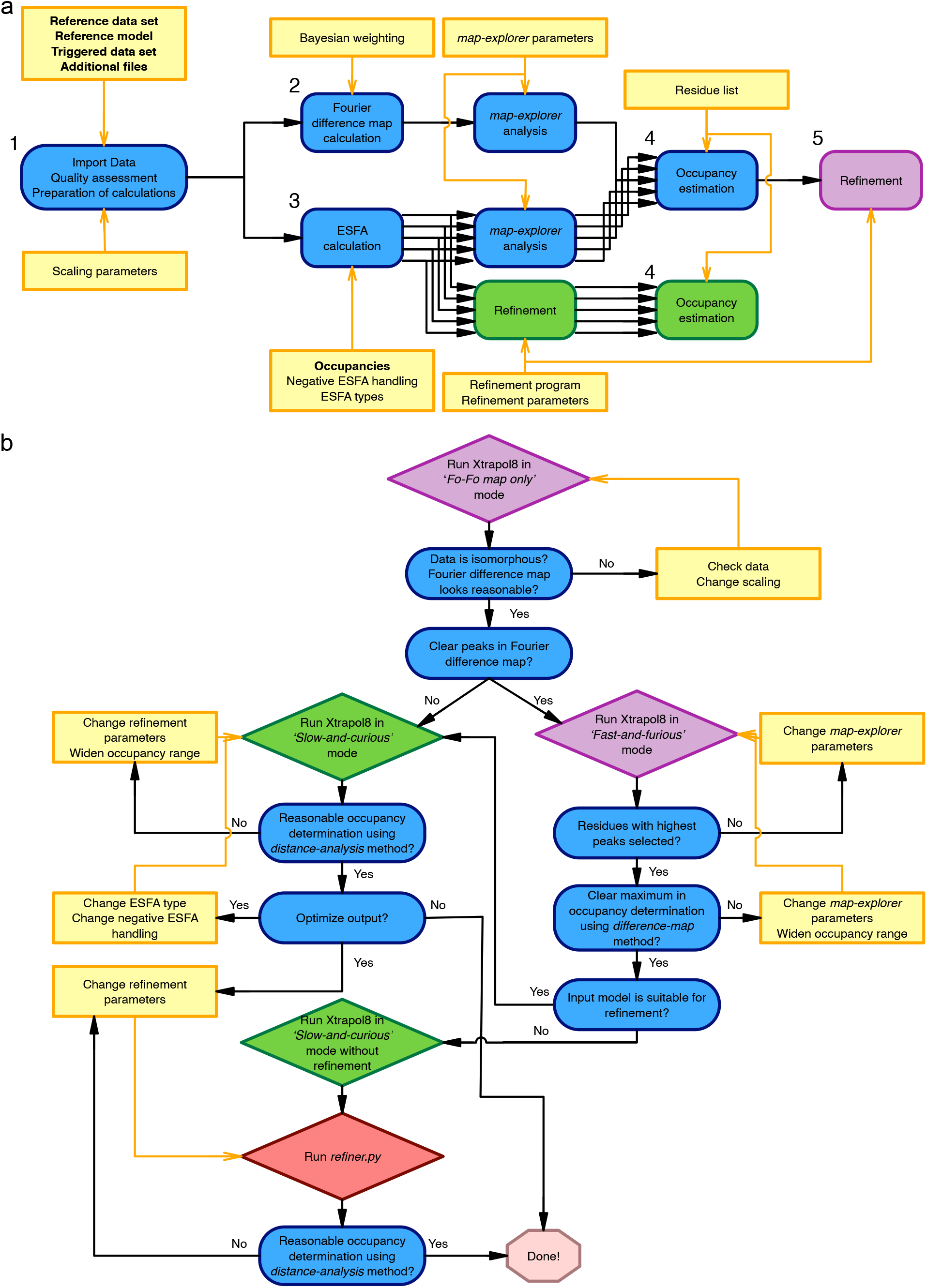
Design and recommended usage of Xtrapol8. **a**, Xtrapol8 roadmap. The four main steps followed by Xtrapol8 are depicted in blue. User input are highlighted by yellow boxes, with obligatory input further highlighted in bold. Steps specific to the ‘*fast-and-furious*’ (default options: *q*-weighting of difference map and ESFAs, rescue of negative ESFAs using the *truncate* method, occupancy determination based on the *difference-map* method; additional step 5: refinement in both reciprocal-space and real-space at the automatically-determined occupancy) and ‘*calm-and-curious*’ modes are boxed in purple and green, respectively. **b**, Suggested workflow for efficient usage of Xtrapol8. Users are advised to first run the program in the ‘*Fo-Fo map only*’ mode in order to evaluate the height of peaks in the difference Fourier map. Secondly, it is recommended to run Xtrapol8 in ‘*fast-and-furious*’ mode (purple boxes) to obtain a crude estimate of the occupancy based on the *difference-map* method, and a first characterization of the triggered state structure. Finer exploration can then be carried out using the ‘*calm-and-curious*’ mode (green boxes), which will refine the occupancy determination based on the *difference-map* method, produce refined structures for all tested occupancies and ESFA strategies, and enable orthogonal occupancy determination by the *distance-analysis* method. Evaluation criteria are shown in blue, user actions are depicted in yellow. See Supplementary methods for a full description.

The green-to-red photoconversion of mEos4b is at the basis of its usage in as a marker in photo-activated localization microscopy (PALM). Yet the existence of reversible dark-states, which form upon excitation of the red-emitting state, has long limited its application in single-particle-tracking (spt) PALM. Based on KX experiments, whereby crystalline mEos4b in the green-emitting state was first slowly converted to the red-emitting state and then illuminated by a 561-nm laser before freeze-trapping, a long-lived dark state was identified whose characterization enabled the design of a new data collection scheme suited for the recording of long tracks in spt-PALM.^9^ In the reference (*red-on*) and triggered (*red-off*) datasets two and three states co-exist, respectively, illustrating how complex investigated structural dynamics may be in TRX and KX studies (Fig. 2 a).

**Fig. 2.**
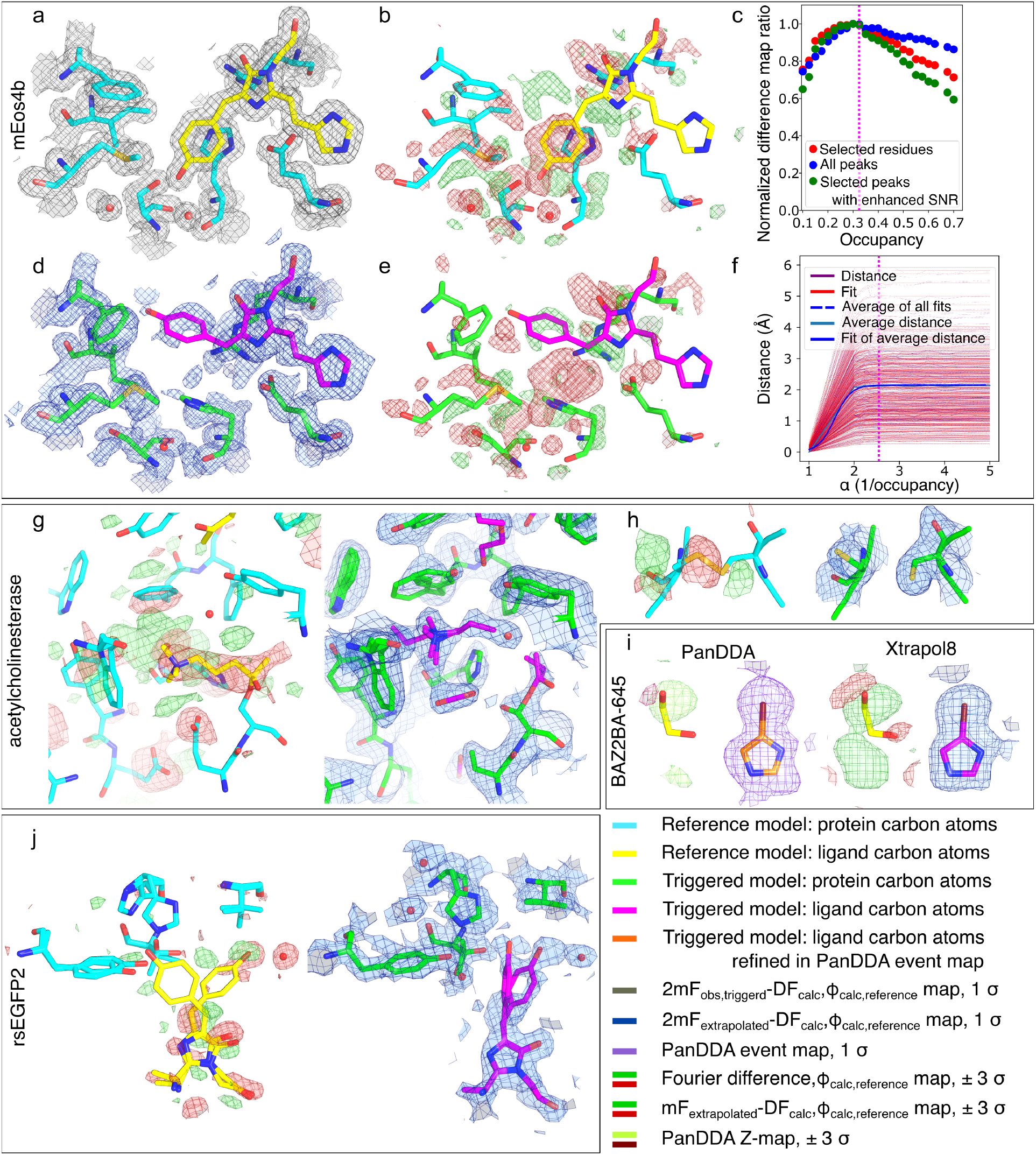
Xtrapol8 enables extraction of low-occupancy states in kinetic, time-resolved and ligand-based crystallography. **a-f**, successful extraction of the mEos4b *red-off* state. The models in cyan and green are mEos4b in the *red-on* and *red-off* state, respectively, as downloaded from the PDB (PDB entry 6GP0 and 6GP1, with the only difference that features of green mEos4b were omitted) with the chromophore indicated in yellow (reference state, **a-b**) and magenta (triggered state, **d-e**). **a**, traditional 2mF_obs_-DF_calc_ electron density after rigid body refinement of the *red-on* state model in the *red-off* data indicates the absence of signal for the *red-off* state. **b**, *q*-weighted Fourier difference map 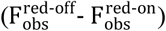 superposed on the *red-on* model. **c**, the *difference-map* analysis method maximizes the information content of extrapolated difference maps and estimates the occupancy to be 0.325 (magenta dashed line). **d-e**, initial *q*-weighted 2mF_extrapolated_-DF_calc_ (**d**) and mF_extrapolated_-DF_calc_ (**e**) electron density maps calculated for an occupancy of 0.325 superposed on the *red-off* model. **f**, the *distance-analysis* method, which approximates the correct occupancy by fitting a sigmoid to the evolution of interatomic distances as a function of all tested occupancies and then retrieves the occupancy value for which 99% of the plateau is reached, returns an occupancy estimate of 0.380. **g-j**, performance of Xtrapol8 on other test cases. In the Supporting results and discussion, we evaluate the versatility and performance of Xtrapol8 by revisiting other time-resolved (TRX), kinetic (KX) and ligand-binding crystallographic studies. For each case, the left panel shows the Fourier difference map superposed on the reference state (with carbon atoms of the proteins and ligands colored in cyan and yellow, respectively), while the right panel shows the extrapolated electron density map superposed on the triggered state model (with carbon atoms of the proteins and ligands colored in green and magenta, respectively). **g-h**, a temperature-dependent KX study was conducted on the covalent complex of acetylcholinesterase with a non-hydrolysable substrate analogue, whereby X-rays were used to radiolytically cleave bonds, including disulfide bridges and the bond tethering the substrate analogue to the catalytic serine.^20^ By use of extrapolation, deeper insights could be obtained revealing two binding poses for the radiolytically produced carbocholine product, trapping of CO_2_ from radiolysis of buried acidic residues (**g**), and breakage of disulfide bridges (**h**). **i**, comparison of the performance of PanDDA and Xtrapol8 in revealing the electron density of a small compound in a fragment-screening study (BAZ2BA-538).^21^ **j**, a TR-SFX study was conducted on the reversibly fluorescent protein rsEGFP2 with aim to determine the structure of the excited state that preludes to isomerization and *off*-to-*on* fluorescence switching.^22^ Xtrapol8 allowed obtaining similar results as published earlier for the 1 ps time delay dataset. In the Supplement we show that Xtrapol8 further allowed extending the results obtained at the 3 ps time delay.

We first ran Xtrapol8 in the ‘*Fo-Fo map only*’ mode, wherein the program stops after calculating a Fourier difference map, to ascertain both isomorphism of the data and occurrence of conformational changes in the triggered dataset (Fig. 1 a, steps 1 and 2). The isomorphism between the reference (PDB entry 6GP0) and triggered dataset (PDB entry 6GP1) is high, with an overall Riso of 0.106 (highest resolution shell Riso= 0.261; 2.5% increase in unit cell volume; Supplementary Fig. 1 a), and the *q*-weighted Fourier difference map 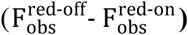 shows strong features on the chromophore and surrounding residues up to a resolution of 1.5 Å (Fig. 2 b). Running Xtrapol8 in the ‘*fast-and-furious*’ mode (Fig. 1 a steps 1-5 with default options, Methods and Supplementary Methods), we tested eight occupancies in the 0.1 to 0.7 range. Occupancy estimation using the *difference-map* method, which is based on the maximization of peak height in the extrapolated difference maps, pointed to the triggered-state (*red-off*) displaying an occupancy between 0.3 to 0.4 (Supplementary Fig. 1 b). Altogether, Fourier difference map calculation, estimation of the occupancy and production of reciprocal and real-space models of the triggered state took 20 min on a mid-range laptop.

Subsequently, Xtrapol8 was run in the ‘*calm-and-curious*’ mode, in which the user can alter various options and full refinement is carried out for each set of user-chosen ESFAs (Fig. 1 a steps 1-4; Methods and Supplementary Methods). By testing 13 occupancies, the final occupancy was determined to be 0.325 (Fig. 2 c) using the *difference-map* method. The initial extrapolated m_Fextrapolated_-DF_calc_ and 2m_Fextrapolated_-DF_calc_ map confirmed the occurrence of large structural changes in the chromophore and surrounding residues (Fig. 2 d and e). Some of these could not be modelled automatically during reciprocal and real-space refinements, requiring manual intervention to model chromophore isomerization and accompanying conformational changes. The final triggered state model was characterized by Rwork/Rfree and CCmask values of 20.91/23.81 % and 91.48 %, respectively. Similar results were obtained when other types of ESFAs and weighting schemes were used (Methods and Supplementary Methods; Supplementary Fig. 2, 3). Only in the case of so-called (*q/k*)Fgenick extrapolated maps,^10^ whereby a direct Fourier synthesis (*i*.*e*. m_ref_|qF_extrapolated_|, φ_ref_) is applied to ESFAs using phases and figures of merit from the dark model, or *k*-weighted extrapolated maps with a high *k*-scale outlier rejection factor, were electron density features less pronounced. This observation suggests that recalculating figures of merit for each set of ESFAs benefits extraction of structural features for the triggered states, enabling to observe structural changes at a lower occupancy. The use of maximum likelihood weighted maps is also likely beneficial, as it allows to take into account not only errors on phases (mref or m) but also those on the measurement and estimation of structure factor amplitudes (D). Specific to mEos4b, similar results were obtained with all possible treatments of negative ESFAs implemented in Xtrapol8, *i*.*e*. when they were rejected, set to 0, replaced by 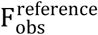 or 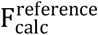, or rescued by use of the *truncate* method (Supplementary Fig. 4). This is likely due to their low amount in ESFAs calculated for an occupancy of 0.325 (ranging from 3.0 to 11.2 % depending on the ESFA calculation strategy). In the Supplementary Results section, we nonetheless show how beneficial proper handling of negative ESFAs can be for the refinement.

The triggered state model was finally subjected to automatic refinement against all sets of ESFAs calculated for different occupancies, enabling to inquire the performance of the *distance-analysis* method for occupancy estimation, which uses the evolution of changes in interatomic distances in the models refined against ESFAs (Methods and Supplementary Methods). To this end we used the *refiner*.*py* script, which allows to relaunch all refinements using another model or refinement strategy and offers the possibility to run the *distance-analysis* method based on the refined models. The occupancy was thereby estimated to be 0.38 (Fig. 2 f), offering orthogonal confirmation for the occupancy determined by the *difference-map* method. The *distance-analysis* method was hardly sensitive to the number of atoms used for the estimation, yielding similar results when either all protein atoms or exclusively atoms with strong difference map peaks were used. Hence, this method could offer solace in cases where the signal-to-noise ratio of the Fourier difference map is low and users can only rely on ESFAs and extrapolated maps.^11^ The introduction of two orthogonal occupancy-determination methods, both based solely on the crystallographic data, hold promise of preventing under-or over-interpretation due to occupancy misestimation.

In the Supplementary Results section, we revisit other TRX, KX and ligand-binding studies that required high-end expertise in crystallography and extensive data-processing, yet could be addressed within hours by use of Xtrapol8 (Fig. 2 g-j). We show in at least two cases (see rsEGFP2 and Shoot-and-Trap test cases) that superior results could have been obtained by the use of Xtrapol8. By enabling automatic elucidation of low occupancy intermediate states, thereby minimizing the time required to extract meaningful results, Xtrapol8 may thus accelerate discoveries in TRX, KX and ligand-binding studies. Unique to Xtrapol8 is that it tackles a variety of issues related to ESFAs, most notably the determination of the triggered state occupancy based solely on X-ray data and the presence of negative SFAs which can result in sub-optimal refinement and electron density maps. Furthermore, Xtrapol8 offers the possibility to calculate all types of ESFAs, which should increase reproducibility while allowing users to make informed decisions as to the method best suited for their project. Lastly, a level of customization is offered on most important parameters, but defaults are carefully set and Xtrapol8 can be run from the command line or via a GUI, so that adequate results are within reach for experienced and novice users.

## Methods summary

Written in python, Xtrapol8 requires only the CCP4^12^ and Phenix^13^ suites to run. Only standard packages and the cctbx toolbox^14^ are used, which are automatically loaded from the phenix.python environment. This facilitates usage as well as transfer between laboratory computers and data collection centers (XFELs, synchrotrons). Xtrapol8 can be run in a ‘*fast-and-furious*’ mode, where the triggered state structure is refined using only the ESFAs calculated at the occupancy determined based on the *difference-map* analysis, or in a ‘*calm-and-curious*’ mode, where refinement is carried out for a range of user-supplied occupancies and the correct occupancy estimated in a later stage (Fig. 1 a). An option is also given to merely calculate Fourier difference maps – the ‘*Fo-Fo map only*’ mode. Two Bayesian-statistics weighting schemes for SFA differences are implemented, referred to as *q*-weighting^5^ and *k*-weighting,^6^ respectively, with the option to apply these either only for the calculation of Fourier difference maps, or for that of ESFAs as well. Specific to *k*-weighting, a rejection factor can be adjusted to vary the influence of outliers on the Fourier difference map and ESFAs. Calculation of ESFAs is carried out by summing the weighted/unweighted (*w*= q/⟨q⟩, *w*= k/⟨k⟩, *w*=1) scaled SFA differences 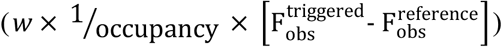 to either observed or calculated reference SFAs (*i*.*e*,. F_additional_ equals 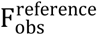 or 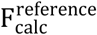, respectively), and the figure of merits used for calculation of initial extrapolated (2m_Fextrapolated_-DF_calc_ and mF_extrapolated_-DF_calc_) maps can either be re-calculated for each set of ESFAs (without phase refinement) or inherited from the reference data. Of important note, the user may choose to test all of these options in a single run of Xtrapol8. Unless specified otherwise, the occupancy of the triggered state is estimated based on the peaks in the m_Fextrapolated_-DF_calc_ electron density map (referred to as the *difference-map* method), and this for each of the requested ESFA calculation strategies. Negative ESFAs, whose amount increases when the occupancy of the triggered state decreases, can be rescued by a variety of methods, the most recommended of which applies French-Wilson^15^ scaling to reconstructed extrapolated intensities, resulting in higher completeness of the data used in map calculations and refinement, and therefore better map quality and refinement statistics (CC_mask_/CC_volume_ and R_work_/R_free_ in the real- and reciprocal-space, respectively). Options are given to refine structures in real-space or reciprocal-space only, or to skip structure refinement. The latter strategy can prove especially useful in cases where the triggered state structure features molecular moieties absent from the reference state structure (*e*.*g*. ligand-binding studies, rapid-mixing TRX studies, pump-probe experiment with caged-compounds) and manual intervention is needed for refinement to converge. Specific to refinement, either phenix.refine^16^ and phenix.real_space_refine^17^ (Phenix) or Refmac5^18^ and Coot^19^ (CCP4) can be used, with options to tweak each program to obtain the best results. Density modification can be carried out to reduce noise levels in the reciprocal-space refined map before real-space refinement is carried out. If the ‘*calm-and-curious*’ mode is selected and structures are refined against all ESFAs calculated for a variety of occupancies, a second estimation of the optimal occupancy can be obtained based on the evolution of structural changes in function of occupancy (*distance-analysis* method), an option particularly useful when Fourier difference maps display low signal to noise ratios. Scripts are provided to relaunch occupancy determinations or refinements with tweaked parameters in the case where the first estimate(s) or refinement results are questioned by the user, or if a better model has become available. Additional details can be found in the Supplementary Methods section. The code and user manual are available at https://github.com/ElkeDeZitter/Xtrapol8.

## Acknowledgements

We thank Kyprianos Hadjjidemetriou, Dominique Bourgeois, Martin Weik, Ilme Schlichting for continuous support and stimulating discussions. IBS acknowledges integration into the Interdisciplinary Research Institute of Grenoble (IRIG, CEA). This work was supported by the Agence Nationale de la Recherche (grants ANR-17-CE11-0018-01 and ANR-2018-CE11-0005-02 to J.-P.C.).

