## Supplementary information for "Xtrapol8: automatic elucidation of low-occupancy intermediate-states in crystallographic studies"

***

### Supplementary information

#### Supplementary Methods

The minimal required input for Xtrapol8 are two data files in mtz or mmcif format, associated to the reference (ground, native, apoprotein) state and triggered (derivative, ligand-bound, intermediate, excited) state, respectively; a model associated to the reference state in pdb or mmcif format; additional files required for proper handling of ligands (such as crystallographic information files), and a list of possible occupancies for the triggered state (Fig. 1 a). The latter is required for the estimation of intermediate state occupancy, which is based solely on crystallographic data and is one of the main features of Xtrapol8. Via complementary methods, such as spectroscopy or binding assays, an estimate of the triggered state occupancy may already be available, requiring to run Xtrapol8 with just a narrow range of possible occupancies. If no estimate is available, a wide range can be tested and the most probable value automatically estimated. It yet remains the user's responsibility to validate and further adjust this estimate, as will be shown by the examples in this manuscript. Depending on the user's experience, many additional parameters can be adjusted including resolution cutoffs, scaling parameters, weighting schemes, and parameters for difference map exploration or structure refinement.

Xtrapol8 takes four main steps when run in the '*slow-and-curious*' mode (Fig. 1 a): 1) Reading of input files and quality assessment, and preparation of the files needed for the next steps; 2) Calculation of the optionally-weighted Fourier difference map, integration of peaks and assignment to reference model atoms; 3) Calculation of extrapolated structure factor amplitudes (ESFAs) using a variety of formalisms (see below) for a user-defined range of possible occupancies, integration of peaks in the extrapolated difference maps and their assignment to reference model atoms, and optional automatic structure refinement; 4) Estimation of the triggered state occupancy and assignment of the most likely extrapolated structure factors and electron density maps for the triggered state structure. The program also has a '*fast-and-furious*' mode in which the refinements in steps 3 are omitted and only performed in a fifth additional step using the extrapolated structure factor amplitudes calculated for the automatically estimated occupancy of the triggered state. In the '*Fo-Fo map only*', only the first and second step are carried out.

#### Graphical User Interface

Xtrapol8 can be controlled through the command line or via a graphical user interface (XtrapolG8) written in *wxPython* as a frontend to the command-line version. The '*Configure'* tab features three input panels arranged according to the different steps (*Input/Output*, *Extrapolation*, *Refinement*) with a selection of options and parameters that are hidden/shown depending on the user-selected expertise level and Xtrapol8 mode (*Fo-Fo map only*, *fast-and-furious*, *calm-and-curious*; Supplementary Fig. 5, 6, 7, and 8). The '*Results'* tab also consists of three panels. The first panel shows the progression of the Xtrapol8 run by tailing the log file. The second panel displays the general output figures, *e.g.* the plots of $R_{iso}$ and $CC_{iso}$ as a function of resolution, the fraction of negative ESFAs as a function of intermediate state occupancy or the results of the occupancy determinations and refinements (Supplementary Fig. GUI_output a). The last panel shows plots specific to a given type of ESFAs and occupancy, accessible through two dropdown menus, *e.g.* the plots of the number of negative ESFAs or ESFA signal-to-noise ratio as a function of resolution. Coot can be directly opened from XtrapolG8 and will load the Fourier difference and extrapolated maps, and refinement output (Supplementary Fig. 9 b and c). For the automatically-determined occupancy, an all-atom distance-difference matrix is also computed between the refined extrapolated-structure and the reference model, whereby the plotted value for each residue corresponds to the sum of the changes in distance for all constitutive atoms.

#### Data handling and quality assessment

The input data files should contain merged data in the form of structure factor amplitudes or intensities. In the case where only the latter are available, they will be converted into structure factor amplitudes using the French-Wilson algorithm[1] using Truncate (CCP4).[2] If both intensities and amplitudes are available, the program will use the amplitudes. Afterwards, Pointless[2] will be run to verify consistent indexing between the triggered and reference datasets, and if needed, will be used to reindex the triggered dataset in respect to the reference dataset. The two datasets are then trimmed so as to retain only reflections present in both sets of observed data and lying within the optional user-defined resolution limits. In the case where no resolution limit is defined, the program will use all data present in the input data files.

Prior to difference map and ESFA calculations, the model-derived calculated structure factor amplitudes are scaled anisotropically to the reference data using mmtbx.fmodel[3,4] while scaling of the triggered to the reference data is performed anisotropically or isotropically by scaleit.[5] A possibility is offered to not scale the observed data at all, to account for the possibility that these may have been scaled already – *e.g.* if merging and scaling of serial time-resolved data was performed using Partialator[6] with the --

custom-split option, or if merging and scaling of rotational triggered data was carried out using the reference dataset in XSCALE[7] or Aimless.[2] At this step, statistical indicators informing on the isomorphism between the input data sets are calculated, which inform users of the probability that both sensible Fourier difference maps and extrapolated structure factor amplitudes can be calculated. Specifically, the R-factor ($R_{iso}$) between the two datasets is calculated using:

$$R_{iso} = \frac{\sum |F_{obs}^{reference} - F_{obs}^{triggered}|}{\sum |F_{obs}^{reference} + F_{obs}^{triggered}|/2} \quad (1)$$

where $F_{obs}^{reference}$ and $F_{obs}^{triggered}$ are the structure factor amplitudes of the reference and triggered states, respectively. An overall $R_{iso}$ value lower than 0.10 indicates strong isomorphism between the two data sets, but values up to 0.25 can still lead to useful difference and extrapolated electron-density maps.[8] The isomorphism-indicating correlation coefficient ($cc_{iso}$) is calculated using:

$$cc_{iso} = \frac{\sum\left[\left(F_{obs}^{reference} - \overline{F_{obs}^{reference}}\right) \times \left(F_{obs}^{triggered} - \overline{F_{obs}^{triggered}}\right)\right]}{\sqrt{\sum\left(F_{obs}^{reference} - \overline{F_{obs}^{reference}}\right)^2 \times \sum\left(F_{obs}^{triggered} - \overline{F_{obs}^{triggered}}\right)^2}} \quad (2)$$

**Experimental Fourier difference maps calculation**

On the assumption that the structural differences between the reference and triggered state(s) are small, *i.e.* only a few specific atoms change positions with no rotation or translation of the molecule in the unit cell, phases of the triggered and the reference states hardly differ, enabling investigators to extract information on the triggered state by calculation of a Fourier difference map:

$$\left|F_{obs}^{triggered} - F_{obs}^{reference}\right|, \varphi_{calc}^{reference} \quad (3)$$

A simple subtraction between the two sets of structure factor amplitudes gives a similar weight to all structure factors, regardless of their measurement precision. Ursby and Bourgeois[9] suggested the use of *q*-weighting whereby a weighting factor is calculated for each difference SFA using Bayesian statistics, based on the magnitude of the difference in structure factors amplitude and the relative errors on their measurements. Thereby unlikely structure factor amplitude differences are downweighed. The improvement of Fourier difference electron density maps by Bayesian weighting was also proposed by Terwilliger and Berendzen[10] who used a similar approach to *q*-weighting, and by Ren *et al.* who introduced a similar but simpler weighting scheme[11] (to which we refer to as *k*-weighting in analogy to *q*-weighting). In Xtrapol8, the user has the choice to calculate a *q*-weighted, *k*-weighted or non-weighted Fourier difference map, but the default is set to *q*-weighting. In the *k*-weighting scheme, the extent to which outliers are downweighed can be adjusted by an additional parameter, that has been set to 0.05[12] or 1.0[11,13] depending on reports, with the first one being the default in Xtrapol8.

During data collection at a synchrotron or XFEL source, it may be that users need to calculate Fourier difference maps to follow the progress of their experiment, while not needing to carry out ESFA calculations and subsequent refinement of extrapolated structures. For such cases, Xtrapol8 can be run in '*Fo-Fo map only*' mode. At the command line, this mode is enabled by setting to "True" the output.generate_fofo_only parameter. When using the graphical user interface XtrapolG8, users need to check the "FoFo only" option in the *Extrapolation* panel of the *Configure* tab.

**Difference map analysis**

An important feature of Xtrapol8 is the *map-explorer* module, which will search for peaks in difference maps, integrate and assign them to the closest reference model atom within a given radius, and to generate an unbiased list of residues whose position changes relative to that in the reference model. Based on this list, the residues with the highest integrated peak volumes can be used to evaluate the occupancy of the triggered state in a later stage of the program. The list can be modified by the user, *e.g.* in cases where only active site or ligand atoms are to be considered for the estimation of the occupancy or if prior knowledge is available as to which atoms undergo significant conformational changes upon triggering of the studied reaction. The stringency in the selection of difference peaks can be adjusted – not only in terms of the maximal and minimal heights for peak selection and integration, respectively, but as well in terms of Z-score filtering. Default values for these parameters are ± 4 σ, ± 3 σ and a Z-score of 2.

**Extrapolated structure factors**

Extrapolated structure factor amplitudes are estimates of the structure factor amplitudes that the triggered state would have produced, had it been present at full occupancy in the crystal. Generally, they can be written as:

$$F_{\text{extrapolated}} = w \times \alpha \times \left(F_{\text{obs}}^{\text{triggered}} - F_{\text{obs}}^{\text{reference}}\right) + F_{\text{additional}} \qquad (4)$$

where α is the reciprocal occupancy of the triggered state (α=1/occupancy) and *w* a potential weighting factor. The weighting factor *w* and the $F_{\text{additional}}$ term have found different interpretations in the literature. Firstly, $F_{\text{additional}}$ was initially assigned to the reference-model derived calculated structure-factor-amplitudes ($F_{\text{calc}}^{\text{reference}}$or $F_{\text{model}}^{\text{reference}}$),[10] but use of measured structure factor amplitudes for the reference state ($F_{\text{obs}}^{\text{reference}}$) was later suggested by Genick and co-workers[14,15] Whether the $F_{\text{obs}}^{\text{reference}}$ should be preferred to $F_{\text{calc}}^{\text{reference}}$ is still under debate,[16] illustrating that it likely depends on the experimental case. Secondly, one can question whether or not the *q/k*-weighting scheme used to reduce the weight of uncertain structure factor differences in the Fourier difference map should also be applied in the calculation of extrapolated structure factor amplitudes (*w* in Supplementary Equation 4),[17–19] leading to yet another nuance. Finally and perhaps more subtly, initial electron density maps are being calculated using the phases of the reference model but the associated figure of merit (*m*) can be chosen to represent the phase agreement between the ESFAs and reference state model, or originate from the reference state.[14,15] Overall, this leads to nine different types of ESFAs and corresponding extrapolated maps (Supplementary Table 1), reflecting the variety of approaches that have been followed depending on the experimental case and laboratory since introduction of extrapolation methods. In Xtrapol8, all types of ESFAs can be calculated at once or only a subset, enabling to compare results obtained by the various approaches. User-selected options for occupancy determination and refinement will apply to all types of ESFAs.

The parameter α is inversely related to the occupancy of the triggered state, but its correct value remains unknown until occupancy determination has been carried out. Two methods are available in Xtrapol8 to estimate the triggered state occupancy, either based on the raw extrapolated maps or on the refined extrapolated models. Hence, extrapolated structure factor amplitudes, maps and models are calculated for each of the user-supplied occupancy values, and occupancy determination is performed at the end of the Xtrapol8 run. By default, and in the *fast-and-furious* mode, only the $qF_{extr}$ type of extrapolated structure factor amplitudes is calculated, but all nine ESFA types – or a subset thereof – can be calculated in a single run in the *calm-and-curious* mode, leading to a maximum of 9 independent occupancy estimations by each of the two occupancy determination methods – or 18 estimates. According to our tests, all should match within a 10% error range.

For each occupancy, ESFAs and extrapolated map coefficients are used in three subsequent refinement steps. First, a reciprocal-space refinement of the reference state input model is carried out against the ESFAs, allowing to account for changes in atomic positions and thereby ameliorate the phases and associated figure of merit *m*, which in turn will result in clearer $mF_{extrapolated}$-$DF_{calc}$ and $2mF_{extrapolated}$-$DF_{calc}$ electron density maps. The latter and the corresponding refined model are further subjected to real-space refinement to account for conformation changes too large to be accounted for in reciprocal-space refinement. In addition, a real-space refinement is also carried out using the reference model and the initial extrapolated electron density map (calculated before reciprocal-space refinement and update of phases and figure of merits), offering an unbiased view of the information content in the extrapolated data and giving the possibility to verify whether sub-optimal refinement leads to phase errors that deteriorate the content of electron density maps. The user has the possibility to perform these reciprocal-space and real-space refinements using either phenix.refine[20] and phenix.real_space_refinement[21] or Refmac[22] and Coot,[23] respectively. An important limitation in streamlining the refinement steps is that they are performed with the input model, which may or may not be adequate to accurately describe the triggered state. With the exception of waters, which can be automatically updated during reciprocal-space refinement when using phenix.refine, no atoms can be removed or added during the Xtrapol8 run. For this reason, it is advisable that automatic refinement is not carried out in studies where covalent bonds are being formed and/or broken or whether the detection and localization of a ligand is of interest.

At the command line, disabling the automatic refinement steps is enabled by setting the refinement.run_refinement parameter to "False". When using the graphical user interface XtrapolG8, users need to un-check the "Perform refinement with" option in the *Refinement* panel of the *Configure* tab. Density modification can aid in ameliorating the phase information and cleaning up the electron density maps, offering a better base for real-space refinement. Regardless of the suites of programs used for reciprocal-space and real-space refinement, density modification is performed using the CCP4 program dm.[5] We indeed found that this program is generally unsensitive to the often non-ideal distribution of ESFAs and their standard deviations.

**Negative extrapolated structure factors**

An often-overlooked issue in structure factor extrapolation is the presence of negative ESFAs. Their origin lies in large negative values of the weighted difference amplitudes (first term in Supplementary Equation 4) that cannot be compensated by the additional amplitude term (second term in Supplementary Equation 4). It should be noted that for a small occupancy, and thus a large α-value, the percentage of negative ESFAs can become large, which will result in decreasing the true completeness, defined here as the completeness of the positive reflections only, below a critical value of 90%. Indeed, negative structure factor amplitudes are not handled by refinement programs so that the refinement is carried out against incomplete data, resulting in weak electron density maps and non-converging refinement. In Xtrapol8, we propose different approaches to correct for this issue. In the first approach, the negative reflections are removed, on the ground that that the error in the measurements of the associated reflections is too large to lead to a reliable estimate.[14] However, rejecting them may strongly alter the resulting extrapolated electron density and furthermore biases the result toward the positive difference amplitudes. In the second approach, the artificial negative reflections are set to zero, under the assumption that true values of these physically impossible negative structure factor amplitudes are weak, but this treatment implies losing important information on their relative strength[1] and again biasing the result toward the positive amplitudes. In the third approach, the negative structure factor amplitudes can be replaced by their corresponding values in either the observed ($F_{obs}^{reference}$) or calculated ($F_{calc}^{reference}$) reference datasets, under the assumption that the strong negative difference amplitude is a consequence of measurement errors in $F_{obs}^{triggered}$ and that the triggered state closely resembles the reference state. Predictably, however, this approach will bias results towards the reference state, which can lead to an unreliable occupancy estimate and alteration of the extrapolated density. Importantly, usage of this option should mirror the ESFA-calculation strategy, *i.e.* negative ESFAs should be replaced with $F_{obs}^{reference}$, in the case where $(q/k)F_{extr}$ and $(q/k)F_{genick}$ ESFAs are used, but with $F_{calc}^{reference}$, when ESFAs are calculated using the $(q/k)F_{extr,calc}$ method. Finally, just as with intensities originating from a diffraction experiment with a fully occupied crystal, also the extrapolated intensities, calculated as the square of the extrapolated structure factor amplitudes multiplied by their initial sign, are supposed to follow the

Wilson distributions.[24] Structure factors can be re-calculated from the extrapolated intensities by the use of an algorithm taking the French-Wilson algorithm[1] into account, *e.g.* Truncate.[2] This approach is that which yields the lowest $R_{work}$ and $R_{free}$ values in reciprocal-space refinement and the most defined electron density around atoms in the model – as judged by higher $CC_{mask}$ and $CC_{volume}$ after reciprocal-space refinement – hence it is the default. This option has yet the drawback that all structure factors are being re-calculated, not just the negative ones. Hence, in cases where negative ESFA amount to less than 5% of all ESFAs, other strategies may become preferable.

**Occupancy estimation**

Determination of the occupancy of the triggered state is one of the main features of Xtrapol8. Indeed, the extrapolated structure factor amplitudes vary as the occupancy changes (Supplementary Equation 4), hence incorrect estimation of the occupancy could lead to under- or over-refinement of structural features in the triggered state. Xtrapol8 contains two complementary methods for the occupancy estimation, both fully based on the X-ray data without any external assumption or prior knowledge about the triggered state, and with only minimal expectations on the behavior of the structure factors and map coefficients. Both methods can make use of all atoms in the asymmetric unit, of residues, ligands and waters which feature the strongest peaks in the Fourier difference map (as identified by *map-explorer*), or of a user-defined selection of residues, ligands and waters.

The first method, the *difference-map* method, relies on the analysis of difference peak heights in the initial extrapolated difference electron density map, *i.e.* that obtained before refinement of the structure ($mF_{extrapolated,occ}$-$DF_{calc}$; last column in Supplementary Table 1). When calculated using a correct reciprocal-space occupancy ($\alpha$) value, this map strongly resembles the Fourier difference map with positive and negative peaks indicating appearing and disappearing features, respectively, as compared to the reference model. Peaks in the initial extrapolated difference electron density map grow in height as the reciprocal occupancy $\alpha$ is increased towards the correct value, and then decrease again due to increase in the map noise level as $\alpha$ further increases. In Xtrapol8, we retain as the correct occupancy that which yields the extrapolated difference electron density map with the highest normalized signal-to-noise ratio. In practice, we first integrate all peaks in the difference map; use the sum of the integrated absolute values of all peaks as a measure of the noise level in the map (N); utilize a Z-scoring approach to single out the most significant peaks and use the sum of the integrated absolute values of these selected peaks as a measure of the signal level in the map (S); plot the normalized S/N ratio in the various initial extrapolated difference electron density maps as a function of $\alpha$, and pick as the correct occupancy that for which S/N is maximized.

The *difference-map* method is most closely related to the experimental data, but it only provides solid results when the Fourier difference map, and thus the extrapolated difference map, contains features that can be clearly distinguished from noise. In cases where this condition is not fulfilled, the peaks in Fourier and initial extrapolated difference map are small and the occupancy determination based on the comparison of the signal-to-noise ratio in the extrapolated difference maps calculated for various occupancies may fail. This might happen when multiple species occupy the same site and reduce each other's signal, if the signal is indiscernible from noise, or if it was not possible to collect enough data for the reference and/or triggered state. In such cases, users should check that the default parameters used for the analysis – *i.e.* maximal and minimal heights for peak selection and integration of ± 4 and ± 3 σ, and a Z-score filtering of 2 – are not too stringent for their data, and re-run the analysis using slightly modified parameters. A stand-alone script, *differencemap_analysis.py*, is provided to re-run the *difference-map* analysis without the need to relaunch Xtrapol8.

If occupancy determination by the *difference-map* method nonetheless fails, users can still rely on the other occupancy determination method, based on the analysis of difference in interatomic distances between the reference model and the models refined against ESFAs and extrapolated maps calculated for an increasing range of α values (*distance-analysis* method). Indeed, we found empirically that upon refinement of the reference model against ESFAs calculated at increasing reciprocal-space occupancy α, atoms that undergo positional changes in the triggered state move from their initial position in the reference model to their final position in the extrapolated structures refined at the correct α and above, with a sigmoidal dependence – *i.e.* after a slow increment, the distance difference increases steadily with α and then levels at a plateau value indicating that only noise is being added to the maps upon further increasing α. This also holds true if the difference signal is too weak to visually standout in the Fourier difference map, but still systematically contributes to the sets of ESFAs, as opposed to noise. Thus, by (1) fitting a sigmoid to the distance changes in function of α for each pair of atoms (*i, j*) ($d_{(i,j)} = f(\alpha)$); (2) extracting $\alpha_{i,j}$ as the α-value closest to the point where those shifts are 99% of the plateau; and (3) plotting the distribution of $\alpha_{i,j}$ and retaining the average $\langle \alpha_{i,j} \rangle$ and maximum in the distribution, an estimate of the α value can be extracted. In order to obtain robust results with this method, the refinements of the compared models should each converge (*i.e.* a sufficient number of refinement cycles should be run so that $R_{work}$ and $R_{free}$ values reach a plateau) and refinement parameters should be chosen carefully. For example, it may be useful to update waters to avoid over-refinement or to change the relative weight of the X-ray and geometry terms during the refinement, which are refinement options in Xtrapol8. By default, the *distance-analysis* method will use atoms to which Fourier difference peaks have been assigned by *map-explorer*, and whose change in distance relative to the reference model is at least 0.05 Å in all extrapolated structures. The program will examine all interatomic distances within the 2-6 Å range (*i.e.* excluding covalent bond and long-range interactions) that can be fit to the logistic function $L/_{1+\,e^{-k(\alpha-\alpha_0)}-1}$ (with L being the maximum value (or 1 under normalized conditions), k the steepness and $\alpha_0$ the α -value of the sigmoidal inflection point) with an $R^2$ of at least 0.95 and a $\chi^2$ of 0.5, and with boundaries set to allow for an occupancy of the triggered state between 1 and the maximum sampled occupancy. In the case that the user wants to alter the list of residues applied to estimate the occupancy, the stand-alone script *distance_analysis.py* can be used to re-run the distance analysis without having to relaunch Xtrapol8. The user can choose to run the analysis using the list of residues featuring peaks in the Fourier difference map before or after Z-scoring, his own list of residues or no list at all in which case all atoms that are present in all models will be examined. Usage of this script is particularly useful in cases where the *refiner.py* script (see below) was used to launch refinements against ESFA and/or extrapolated maps based on a manually modified model. Note that in the ideal case where strong peaks are visible in the Fourier difference map, the *distance-analysis* method can be used as an orthogonal occupancy determination method to confirm the occupancy estimation by the *difference-map* method.

##### *'fast-and-furious'* versus '*calm-and-curious'*

In the default '*calm-and-curious'* mode, Xtrapol8 performs calculations in the most thorough way possible. For rapid, on-(beam)line evaluation of data content, or to estimate the parameters for a next run, calculations can be expedited using the '*fast-and-furious'* mode. In this mode, default parameters will be used for most arguments, *i.e.* only the *q*-weighted Fourier difference map and ESFAs and map coefficients of the $qF_{extr}$ type will be calculated, and refinements will only be performed using the structure factors and maps associated with the occupancy deemed most probable based on the *difference-map* method (Fig. 1 a, steps 1-5). As a consequence, occupancy determination using the *distance-analysis* method is not possible in the '*fast-and-furious'* mode. As most of the Xtrapol8 runtime is occupied by the refinement steps, this mode reduces the overall runtime significantly, allowing crystallographers in the midst of an experiment to take decisions swiftly.

##### Output

The output of an Xtrapol8 run is stored in an output directory whose name can either be defined by the user or automatically (in which case it is called 'Xtrapol8'). This directory will contain the Fourier difference map, a log file, a Pymol session, and various figures meant to help in the evaluation of data quality and interpretation of results (Supplementary Fig. 10). The log file contains the most important information, such as statistics on the input files, the location of output files, the number of negative ESFAs and the results of the occupancy estimation. A Pymol script is written that enables loading the maps and models for the various occupancies, assigned into a Pymol state object so that the results of the full extrapolation can be visualized as a movie starting with the reference state model and maps and progressing towards the triggered state models and maps extrapolated for decreasing occupancies (*i.e.* increasing α). A subdirectory is created for each tested occupancy value, with the name indicative of whether or not ESFAs were *q/k*-weighted. This subdirectory will contain the ESFAs as well as the results from refinements and coot sessions for the automatically determined occupancy.

**Refiner.py**

In the case where the triggered state structure features different atoms as compared to the reference state (ligand based studies, rapid-mixing TRX studies, pump-probe TRX studies with caged-compounds; see the cases of BAZ2BA-x538 and Shoot-and-Trap below), or if the triggered state diverges so much from the reference state that neither reciprocal-space nor real-space refinement can model the structure (*e.g.*, large conformational changes such as fluorescence protein chromophore isomerization; see the case of mEos4b), a better model will become available for refinement of the triggered state upon manual intervention. The latter may then be subjected to reciprocal-space and real-space refinement against all ESFAs pre-calculated for a range of occupancies using the stand-alone *refiner.py* script distributed with Xtrapol8. The script, which at present only works with phenix.refine[20] and phenix.real_space_refine,[21] can take lists of specific arguments for the real and reciprocal-space refinement, provided by the user in the form of Phenix input files (we refer to the documentation of Phenix[25] concerning supported input files and parameters). The *refiner.py* script offers the option to recalculate the occupancy by applying the *distance-analysis* method to the refined models. A list of residues can be provided (*e.g.*, the list written by Xtrapol8 based on the analysis of the Fourier difference map either before or after Z-scoring) but in the case none is given the analysis is performed using all atoms in the model. A compulsory input is the Xtrapol8_out.phil file, written at the end of the preceding Xtrapol8 run, and it is advised to run the *refiner.py* script in the same directory as that used to launch Xtrapol8. Just as for Xtrapol8, *refiner.py* should be launched with phenix.python.

**Good users' practice**

The main ambivalence with extrapolation is that in order to calculate the most plausible set of ESFAs, the occupancy must be known. However, occupancy estimation based on X-ray data can only be carried out after the initial extrapolated electron density maps (*difference-map* method) and/or refined extrapolated models (*distance-analysis* method) have been produced. Therefore, the scheme outlined in Fig. 1 b is advised to obtain optimal results in the shortest time. Running Xtrapol8 in the '*calm-and-curious*' mode with improper settings would indeed result in a waste of (computation) time, notably when the program is used to monitor the progress and success of an experiment. Best thus appears that users run Xtrapol8 in the '*Fo-Fo only*' or '*fast-and-furious*' mode until good parameters are found (Fig. 1 b). A first good sign is that scaling statistics look reasonable, *i.e.* $R_{iso}$ and $CC_{iso}$ display decent values ($R_{iso} < 0.20$ ; $CC_{iso} > 0.80$) even in the highest resolution shell ($R_{iso} < 0.40$; $CC_{iso} > 0.60$). In the case they are not, users should check their data processing and merging statistics, and the unit cell parameters of the reference and triggered datasets, to increase their chances of getting useful results.

Ideally, the Fourier difference map should cover specific residues and hint to conformational changes in accordance with any prior knowledge, *e.g.* peaks covering active sites residues, and the $2mF_{extrapolated}$-$DF_{calc}$ electron density maps should be of acceptable quality, *i.e.* cover all atoms in the model without breaks in the electron density nor spurious peaks in the solvent channels. Specific to serial crystallography, and provided that difference peaks are visible in the Fourier difference maps, users should continue collection of triggered and reference data until the peak height does not increase anymore as a function of the number of indexed patterns (using the '*Fo-Fo only*' option to stop Xtrapol8 after the generation and analysis of the Fourier difference map). The authors note that they have witnessed cases where rapid calculation of Fourier difference and extrapolated electron density maps were the means by which problems in the experimental setup or crystalline system were diagnosed during TRX experiments. Such cases illustrate the usefulness of having at hand a program like Xtrapol8 in the course of a TRX or KX experiment. It may be, however, that no peaks are visible in the Fourier difference maps, in which case users should proceed with extrapolation so as to verify whether or not information is present in the $2mF_{extrapolated}$-$DF_{calc}$ and $mF_{extrapolated}$-$DF_{calc}$ electron density maps. When this is not the case, the scaling parameters and resolution boundaries may need to be altered.

We note that in order to get a decent estimate of the occupancy, it may be needed to set and optimize a suitable strategy. If features in the Fourier difference map are clear, the default *difference-map* method with default parameters should provide proper results for the estimated occupancy. If nonetheless occupancy determination is poor, a wider range of possible occupancies should be tried. To increase the stringency of the *difference-map* method, users may need to decrease or increase the number of difference peaks used in the *map-explorer* analysis, which can be done by tweaking (1) the maximum (*peak* parameter; default is $\pm 4\ \sigma$) and minimum height (*threshold* parameter; default is $\pm 3\ \sigma$) for peak selection and integration, respectively; (2) the Z-score filter applied to integrated peaks to select the highest ones for occupancy determination (default is 2, corresponding to the ~4.5% highest peaks); and (3) the search radius for assignment of a peak to an atom (default is set to the maximum resolution). When the features in the Fourier difference map are weak, the *difference-map* analysis will not allow to discriminate between tested occupancy values, in which case the *distance-analysis* method can offer solace. This method however relies on the real-space refined models, and thus requires Xtrapol8 to be run in '*calm-and-curious*' mode. Alternatively, users that have already run Xtrapol8 in the '*fast-and-furious*' mode can use the *refiner.py* script to produce reciprocal-space and/or real-space models refined against ESFA and extrapolated maps, respectively. The script offers the possibility to carry out the occupancy estimation using on the *distance-analysis* method at the end of the refinement steps.

It is evident that not only difference signals but as well noise levels increase upon extrapolation of difference data. An illustration of this can be seen from the shape of the occupancy-determination plot produced by the *difference-map* method, which most often shows a maximum rather than a plateau, and frequently needs human interpretation. It may yet be that the occupancy of the triggered state is low, and that the only means by which the triggered state structure can be revealed is by coping with the high noise level in ESFAs and extrapolated maps. In such cases, refinement parameters have to be carefully validated. It may be advisable to reassess the validity and number of water molecules used in refinement, which Xtrapol8 offers to do using the ordered_solvent option in phenix.refine. By this means, the $R_{work}/R_{free}$ values can be improved by a few percent, which in turn results in better maps. In the case where $R_{work}/R_{free}$ values do not converge during refinement, the user may opt for increasing geometry restraints, at the price however of reducing the capacity of atoms to change position at each cycle, resulting in the need for more refinement cycles. Options to control the weight of the X-ray/geometry terms in the maximum likelihood refinement target are included in Xtrapol8, which directly bind to the relevant options in phenix.refine or refmac5. Specific to the latter, a jelly-body refinement mode exists, access to which is also implemented in Xtrapol8. Another option to cope with high noise levels in extrapolated maps is to use density modification under the reasonable assumption that even in the case of extrapolated data, solvent and protein regions are distinct, the electron density histogram is preserved, and phases can be extended in multiple resolution steps. In the case where density modification is performed, the real-space refinement steps will be carried out using the density-modified reciprocal-space refined maps. In cases where $R_{work}/R_{free}$ are too high to achieve trustable refinement (> 50%), *i.e.* melioration of phases and figures of merit, users may fully skip reciprocal-space refinement (by setting the number of reciprocal refinement cycles to 0), and solely perform real-space refinement in the initial extrapolated maps. If this strategy is followed and density modification selected, the two real-space models produced by Xtrapol8 are based on the initial and density modified extrapolated maps, respectively. Otherwise, the two real-space models are based on the initial and refined $2mF_{extrapolated}$-$DF_{calc}$ extrapolated maps. Of note, in the more favorable case where structures can be refined properly and $R_{work}/R_{free}$ values both decrease and converge, common refinement strategies are advised. Hence, all important options in phenix.refine (including simulated annealing, occupancy refinement, individual and grouped B-factor refinement (adp), tls refinement, map-sharpening, etc.) and refmac5 (twinning, overall and isotropic B-factor refinement, tls refinement, map-sharpening, etc.) have been integrated in Xtrapol8. It must be noted that in our tests, refinement using Refmac5 did not converge in cases where the triggered state occupancies were lower than ~20%, leading to errors in phases and degradation of the extrapolated maps and models.

**Framework of the results presented in this paper**

We applied Xtrapol8 on several sets of published TRX, KX and ligand-binding data. These examples were run using Phenix[25] v.1.19 and the associated cctbx modules,[26] CCP4[5] v.7.1 and Coot 0.8.9 or 0.9.4, unless stated differently. Figures were prepared using Pymol 2.5.[27]

#### Supplementary Results

##### mEos4b additional results

The structure factors of mEos4b in the *red-on* and *red-off* states were downloaded from the Protein Data Bank (PDB entry: 6GP0 and 6GP1, respectively). The cif file for the chromophore (3-letter code IEY for the red state chromophore) and the altered phenylalanine (3-letter code NFA; a consequence of green-to-red photoconversion) were generated using phenix.elbow. In these data, it is notable that because of incomplete green-to-red photoconversion, both the *red-on* and *red-off* states contain a ~ 50 % contribution from the non-photoconverted green state. This implies that the two data set only differ by the contribution of the *red-off* (triggered) and *red-on* (reference) state. Therefore, we decided for this case study, to simplify the reference input model by removing the green state structural features and setting the red-state features to an occupancy of 1.0 for its usage in Xtrapol8. To evaluate the performance and complementarity between different strategies offered by Xtrapol8, we tested: i) the benefit of Bayesian weighting of difference SFAs on Fourier difference map quality, ii) the accomplishment of the different types of ESFAs (Supplementary Table 1) in enabling accurate modelling of the triggered state, iii) the pertinence of Bayesian weighting of difference SFAs prior to calculation of ESFAs , and iv) the opportune use of our different approaches to handle negative ESFAs.

Xtrapol8 enables calculation of three types of Fourier difference maps, which differs in the weighting scheme applied to difference SFA, *i.e.* no weighting, *q*-weighting or *k*-weighting. Specific to the latter, we further evaluated the effect of varying the additional user-modifiable scaling parameter to downweigh outliers during *k*-weighting. Indeed, in the literature, values of 0.05 to 1.0 have been used.[11,12] We found that in the mEos4b case, the *q*-weighted map and the *k*-weighted maps calculated with a *k*-weight scale factor of 0 - 0.3 are those featuring the highest quality, as judged from the height of peaks revealing presence and disappearance of the *red-off* and *red-on* states, respectively, and from the reduced number of spurious peaks attributable to noise (Supplementary Fig. 2).

We next challenged the different types of ESFAs in terms of the quality of the $2mF_{extrapolated}$-$DF_{calc}$ electron density map. Examination of extrapolated maps calculated at with an occupancy of 0.325 (Supplementary Fig. 3) point to damped electron-density features for the triggered (*red-off*) state in $2mF_{extrapolated}$-$DF_{calc}$ maps calculated using the (*q/k*)Fgenick type of ESFA [14,15] and, accordingly, electron density remains fully visible for the initial reference (*red-on*) state. Remarkably, a similar trend is observed upon increasing the *k*-weighting outlier scale factor, which suggest that the result will tend towards the reference state if large structure factor differences are rejected from the map-calculation. It should be noticed that while informative Fourier difference maps can be calculated up to a *k*-weight scale factor of 0.3, the bias towards the reference manifests itself as of a *k*-weight scale factor of 0.1 – *i.e.*, only *k*-weight scale factors as low as 0 or 0.05 appear acceptable for the calculation of ESFAs in this case. As noted by the authors in the initial publication, the chromophore in the *red-off* state most probably contains multiple conformations to which a main population can be modeled. This can explain why CCmask values lie in the 0.5 – 0.6 range and the *red-off* chromophore not being entirely covered by clear electron density. Specific to the usability of the various types of ESFAs to estimate the occupancy based on the *difference-map* analysis, we found the predicted occupancy to lie between 0.3 and 0.4 in most cases.

Last we tested the effect on extrapolated map quality and triggered state structure refinement of the various handling schemes implemented in Xtrapol8 to cope with negative ESFAs. For this comparison, we reasoned that the handling of negative ESFAs should mirror that of missing reflections for the calculation of $2mF_{extrapolated}$-$DF_{calc}$ electron density maps, as taken care of by mmtbx.map_tools.[26] Thus, automatic filling of missing map coefficients of the $2mF_{extrapolated}$-$DF_{calc}$ map was performed when the truncate-, $F_{obs}^{reference}$-, $F_{calc}^{reference}$- and zero-strategies were used, but not when the reject-strategy was applied. In the specific case of mEos4b, with a low fraction of negative ESFAs (2.5 – 10.2 % depending on the ESFA type), the weighting scheme and type of extrapolation appears to have a higher influence on the extrapolated electron density and refinement R-values than the negative ESFA strategy (Supplementary Fig. 3, 4 and 11).

**Identification of ps-lived excited-states in a reversibly switchable fluorescent protein (rsEGFP2) by use of time-resolved serial femtosecond crystallography**

rsEGFP2 is a reversibly switchable fluorescent protein that can be toggled back and forth between a fluorescent *on*-state and a non-fluorescent *off*-state by subsequent illumination with cyan and ultra-violet light, respectively. Just like in mEos4b, the chromophore is a conjugated system consisting of an aromatic phenol ring and an imidazolinone ring that are maintained coplanar by a methylene bridge. The chromophore features a *cis* anionic configuration in the fluorescent *on*-state, but is *trans* protonated in the non-fluorescent *off*-state. Structures of these states have been solved by rotation-based cryo-crystallography at 100 K (PDB entries 5DTX and 5DTY for the *on*- and *off*-state structures, respectively)[28] and by serial crystallography at room-temperature (PDB entries 5O89 and 5O8A for the *on*- and *off*-state structures, respectively).[18] Long debated has been whether the first step of photoswitching is isomerisation or protonation, and the question was addressed experimentally by performing time-resolved serial-femtosecond crystallography (TR-SFX) at an XFEL[18] on a slurry of rsEGFP2 microcrystals. *Off*-to-*on* photoswitching was followed after pulsed laser illumination at 400 nm to pump the system to the *on*-state. The crystalline system was probed at time delays $\Delta t$ of 1 and 3 ps after the 400-nm light trigger, with aim to shed light on the structure and decay of the excited state. As a reference, data was also collected on untriggered crystals (*i.e.* crystals in the initial *off*-state), enabling calculation of a Fourier difference map for the two-time delays ($F_{obs}^{laser\text{-}ON\text{-}1ps} - F_{obs}^{laser\text{-}OFF}$ and $F_{obs}^{laser\text{-}ON\text{-}3ps} - F_{obs}^{laser\text{-}OFF}$). For the 1 ps time delay, ESFAs were calculated and the occupancy of the triggered state was determined to be 7.5 %, based on a forerunner of the *difference-map* method applied to manually-selected residues. This state was found to consist of two conformers; a twisted intermediate (referred to as model T in Ref[18]), with a quasi-perpendicular positioning of the phenol and imidazolinone groups in a so-called 'twisted' chromophore; and a *trans*-like intermediate, with the ring moieties very close to those of the *trans* protonated *off*-state (referred to as model P in in Ref[18]). Modelling of these intermediates was complicated by the comparatively poor quality of the extrapolated maps, due to low occupancy of the triggered state and to a high amount of negative ESFAs, which translated to a reduced completeness of the data used in refinement and map calculations. For these reasons, the authors then refrained from calculating ESFAs from the data collected at Δt=3 ps, which were of lower quality than those collected at the Δt=1 ps time delay. Hence, these pump-probe TRX data collected on rsEGFP2 appeared to us as an obvious test case for Xtrapol8.

The deposited structure features two full chains, present as alternate conformers, reflecting the mixture of *on*- and *off*-states (10 and 90 %, respectively) present in rsEGFP2 crystals upon pre-illumination at 488 nm (PDB entry 5O8A). To simplify the model used for phase calculation, and avoid possible bias from the reference structure, we performed a novel refinement of the *off*-state structure (laser-OFF dataset, 1.7 Å resolution) whereby instead of a full-length *on*-state conformer at an occupancy of 10%, we here accounted for the residual *on*-state by addition of alternate conformers only at loci featuring peaks in the $mF_{obs}$-$DF_{calc}$ maps (Supplementary Fig. 12 first column). The $R_{work}/R_{free}$ values of the two reference models are similar, viz 13.4 / 17.5 % and 14.7 / 17.6 % for the re-refined and deposited models, respectively. Using the laser-OFF (reference) and laser-ON-1ps (triggered) datasets, and the new reference model, we ran Xtrapol8 in the '*Fo-Fo map only*' mode to generate *q*-weighted and *k*-weighted Fourier difference maps, and then in the '*calm-and-curious*' mode to compute unweighted, *q*-weighted and *k*-weighted ESFA for ten evenly-spaced occupancies between 0.05 and 0.7 (Supplementary Fig. 13). Initial occupancy of the triggered state was determined using the *difference-map* method, and amounted to 0.075-0.10, depending on whether *q*-weighting (0.075), *k*-weighing (0.10) or no weighting (0.10) was used. In the *'calm-and-curious'* mode, automatic refinement was performed at all occupancies. We found *k*-weighting of ESFAs to be superior to *q*-weighting, yielding 2 - 8 % improvement in initial $R_{free}$ depending on occupancy. With *k*-weighting, and regardless of the occupancy at which extrapolation is carried out, the standard deviation on the ESFAs (σ(kFextr)) hardly varies with resolution (Supplementary Fig. 13 c). Contrastingly, the standard deviations on unweighted (σ(Fextr)) and *q*-weighted (σ(qFextr)) ESFAs show a similar trend, being surprisingly high at low resolution, minimal around 2.5 Å, and rising again at high resolution. Probably, this difference in the determination of σ(ESFA) is at the root of the higher $R_{work}$ and $R_{free}$ values observed for the unweighted and *q*-weighted extrapolated structures, as compared to their *k*-weighted counterparts. The plots showing the weighting factor in function of resolution (Supplementary Fig. 13 d) reveal that lower *k*-weights are at the origin of the reduced σ(ESFA) on k-weighted ESFAs at low and high resolution. Thus, *k*-weighting is useful to account for measurements errors at low resolution, here due to the hollow nature of XFEL detector, and at high resolution, due to reduced diffracted intensities. In the refined extrapolated maps, density was visible for the P and T states in the $2mF_{extrapolated}$-$DF_{calc}$ map, regardless of the method to treat negative ESFAs and weighting (Supplementary Fig. 14), and they could be modelled manually in Coot. The density for model T is best defined in the *q*-weighted maps, despite an overall lower $CC_{mask}$ compared to *k*-weighted maps; possibly, this is due to a damping of structural differences in the *k*-weighted ESFA, which results in reference state features being more present (and inversely for the triggered state features) in the corresponding maps, thus resulting in a better-defined electron density and higher $CC_{mask}$. Focusing on the *q*-weighted maps, the extrapolated map calculated from ESFA generated using the truncate method is that which displays the best overall quality, as judged from $CC_{mask}$. After inspection of the rest of the protein structure and of waters for the structure calculated using *k*-weighted ESFAs at an occupancy of 0.1, the resulting model was subjected to automatic reciprocal-space and real-space refinement against all extrapolated datasets and maps using the *refiner.py* script, offering a view of how conformation and occupancy of the two intermediate states are influenced by the extrapolation occupancy (Supplementary Fig. 15, 12 and 16), and affording to obtain another estimate of the correct value of the occupancy by the *distance-analysis* method (Supplementary Fig. 17). The distance-refinement method fairly reproduces the occupancy determination from the *difference-map* method, regardless of whether *q*-weighted or *k*-weighted or unweighted extrapolated maps are used for real-space fitting, or of whether all atoms or only atoms with $F_{obs}^{laser\text{-}ON} - F_{obs}^{laser\text{-}OFF}$ peaks or only atoms with the highest $F_{obs}^{laser\text{-}ON} - F_{obs}^{laser\text{-}OFF}$ peaks are considered for the estimation. Hence, the *distance-analysis* method can be used to judge extrapolation results and determine the triggered state occupancy even in cases where $F_{obs}^{laser\text{-}ON} - F_{obs}^{laser\text{-}OFF}$ peaks are weak or absent. This may be the case when conformational changes are small in amplitudes or when the triggered state consists (as is the case here) of a mixture of close conformers whose positive and negative density peaks $F_{obs}^{laser\text{-}ON} - F_{obs}^{laser\text{-}OFF}$ overlay resulting in a reduction of the observed difference electron density for each. When all atoms are used, hints can be obtained regarding the presence of different intermediate states present at different occupancies in the triggered state dataset, *e.g.,* the P and T models. Refinement results obtained using the *q*-weighted and *k*-weighted ESFA were similar, and point to an occupancy of 10-15%, in reasonable agreement with the *difference-map* method (Supplementary Fig. 16).

Encouraged by these results, we asked whether or not refinement could also be performed against ESFAs calculated for the 3 ps time delay. At the time of publication, the authors refrained from carrying out refinement against ESFAs because of the large amount of negative ESFAs (~20% at 20% occupancy) that resulted in high $R_{work}$ and $R_{free}$ values. We ran Xtrapol8 in the *'calm-and-curious'* mode with ten evenly-spaced occupancies between 0.05 and 0.7. We used the *difference-map* method to estimate the occupancy of the triggered state(s), which returned values of 0.05, 0.075 and 0.1 when *k*-weighting, *q*-weighting or no weighting was used, respectively (Supplementary Fig. 18). Reciprocal-space and real-space refinement were carried out using the same protocol as above. From the extrapolated *k*-weighted $2mF_{extrapolated}$-$DF_{calc}$ map, we were able to model two chromophore conformers, viz. a pseudo-*trans* conformer and a pseudo-*cis* conformer Supplementary Fig. 19 and Supplementary Fig. 12). This model was refined against all sets of EFSA using the refiner.py script again pointing to an occupancy to 5 – 10 % in line with the prediction of the *difference-map* method (Supplementary Fig. 20).

**Xtrapol8 performance in ligand binding and comparison with PanDDA**

Fourier difference maps and structure factor extrapolation can also prove useful in cases where the complex structure of a protein with a poorly-occupied ligand is of interest. Indeed, in such cases as well, low occupancy translates to poorly-defined electron density making identification of the binding pose challenging if not impossible. To date, the best solution for such cases is the real-space subtraction and map-deconvolution approach implemented in the software PanDDA,[29] whereby a reference electron density map, averaged from hundreds of ligand-free datasets, is for each ligand subtracted from the electron density map obtained after automatic refinement. To compare the performance of Xtrapol8 and PanDDA in their usability for fragment screening, we selected two datasets, BAZ2BA-x645 (reference) and BAZ2BA-x538 (triggered), from the BAZ2BA fragment screening dataset (downloaded from Zenodo[30]) that was used to showcase the performance of PanDDA in its original publication.[29] Interestingly, the BAZ2BA-x645 dataset was also collected from a crystal soaked with a ligand, but as the latter did not bound, the dataset was automatically identified as a reference dataset by PanDDA. In a regular Xtrapol8 experiment, the reference data set would ideally emanate from an unsoaked crystal. Whereas only 2 data sets from the BAZ2BA set were used to run Xtrapol8, we ran PanDDA,[29] as part of ccp4-7.0, on the complete set, consisting in hundreds of datasets, without defining a resolution cutoff.

We first ran Xtrapol8 in the '*Fo-Fo only*' mode. The overall $R_{iso}$ value between the BAZ2BA-x645 and BAZ2BA-x538 datasets is 10.2 % over the full resolution range (28.9 - 1.76 Å), but raises steeply to 30.6% in the highest resolution shell. Therefore, data were cut to 1.85 Å, leading to an overall $R_{iso}$ of 9.8 % and high resolution $R_{iso}$ of 23.9 % (Supplementary Fig. 21 a). Xtrapol8 was subsequently run in the '*fast-and-furious*' mode, testing 10 occupancy values between 0.05 and 0.5. As in ligand-binding studies the reference model does not contain the modelled ligand, it is advisable that no refinement steps are run after extrapolation. The *difference-map* method pointed to an occupancy between 0.20 to 0.30. Xtrapol8 was thereafter rerun in the *'calm-and-curious'* mode, using the truncate method to handle the negative ESFAs. Ten evenly-space occupancy values between 0.15 and 0.33 were tested, each using various strategies for weighting and ESFA-calculation. Because refinement was not carried out, occupancy estimation could only be performed automatically using the *difference-map* method. With ESFAs calculated using the *q*Fextr method, we found an occupancy of 0.25 for the 4-bromoimidazole ligand-bound state (Supplementary Fig. 21 b). Similar occupancies were predicted when other types of ESFAs were used, ranging from 0.17 (*k*Fextr_calc) to 0.27 (*k*Fgenick), or when PanDDA was used

(0.24). The extrapolated *q*-weighted $2mF_{extrapolated}$-$DF_{calc}$ electron density map produced by Xtrapol8 is similar to the event-map calculated by PanDDA for the brominated ligand (Supplementary Fig. 21 c and d). Yet, the Fourier difference map of Xtrapol8 surpasses the z-map of PanDDA in that not only the bromide, but as well the imidazole moiety, are visible at a sigma-level of 4 σ (Supplementary Fig. 21 e and f). Fourier difference peaks could also be found on the protein (Supplementary Fig. 21 g and h), pointing to the conformational changes undergone by the protein upon binding of the ligand.

The BAZ2BA test-case provide a good example for the use of Xtrapol8 in structural enzymology where, alike in this example, the reference and triggered state may contain different ligands. This may also apply in the case of time-resolved experiments on crystalline proteins complexed with caged-compounds in which the compound breaks in multiple moieties upon reaction triggering and might not bind to the protein prior to uncaging. In such cases, it may only become useful to perform occupancy determination using the *distance-analysis* method after the triggered structure has been (manually) refined using an occupancy estimate from the *difference-map* method (best case scenario) or decided upon based on the visual inspection of extrapolated maps. A user could then decide to use the *refiner.py* script to refine the manually-fitted triggered-state model against all sets of ESFAs calculated for the various occupancies. This would then open the door to orthogonal occupancy determination by the *distance-analysis* method.

**Shoot-and-Trap study of a complex of acetylcholinesterase with a non-hydrolysable substrate analogue**

By enabling the rapid hydrolysis of the neurotransmitter acetylcholine into acetate and choline, acetylcholinesterase (AChE) terminates impulse transmission at cholinergic synapses, restoring neuronal excitability.[31] Such synapses are present in the central nervous system, where they support cognitive functions, as well as in the parasympathetic branch of the peripheral autonomous nervous system.[31] Therefore the enzyme is essential, as well as the requirement that it functions at a rate that is nearly diffusion-limited, *i.e.* ~1000-10000 $s^{-1}$ depending on species. Because the active site of AChE is buried at the bottom of a ~20 Å deep aromatic gorge,[32] the molecular basis for the high turnover rate long remained unclear, and likewise for the mechanism underlying substrate inhibition at high concentrations.[33] It was proposed based on early molecular dynamics simulations that the choline product could exit through a backdoor,[34] which would transiently open at the bottom of the gorge, where the enzyme wall is the thinnest, following rupture of H-bonds between the polar side chain atoms of W84, W432 and Y442. Thereby, substrate and product traffic in the active site gorge would be alleviated, contributing to the high turnover rate. In 2006, structures of AChE in complex with isosteric hydrolysable and non-hydrolysable substrate analogues were produced, revealing the molecular basis for substrate binding and inhibition.[35,36] The non-hydrolysable substrate analogue (4-oxo-N,N,N-trimethylpen-tanaminium; OTMA) can bind both at the active and the peripheral site, located at the bottom and entrance of the gorge, respectively, leading to closure of the gorge at mid height. In the active site, the analogue covalently binds to catalytic Ser200 from the catalytic triad, with the H-bonding pattern between the catalytic triad residues (E327, H440 and S200) remaining unaffected. However, the electron density of the analogue was reduced around the C5-C6 bond, which replaces the hydrolysable C-O bond of the natural substrate, illustrating the electron-withdrawing effect of covalent binding to catalytic S200. It is this observation that led to a subsequent study on this complex, whereby data was collected before and after rupture of the substrate analogue C5-C6 bond by prolonged exposure to X-rays, in an approach coined "Shoot-and-Trap".[37] By means of *q*-weighted Fourier difference map calculations, it was shown that the carbocholine radiolysis product reorients in the active site following radiolytic cleavage of the substrate analogue at 100 K, but escapes the active site at 150K. A succession of positive and negative peaks in the map were observed on the residues proposed to be involved in the opening of a putative backdoor, offering first structural support for its existence.[34] Disulfide bond disruption upon exposure to X-rays was also evident from the *q*-weighted Fourier difference maps at both temperatures. Additionally, evidence was obtained for the trapping two $CO_2$ molecules produced by radiation-induced decarboxylation of buried acidic residues at 100K.

At the time, however, structure factor extrapolation and reciprocal-space refinement were not attempted by the authors; hence we wondered if further insights could have been obtained by use of extrapolation methods. We therefore subjected the 100 K reference (~3 MGy, *i.e.* $1/10^{th}$ of the experimentally determined Garman limit[38]) and triggered (~9 MGy, *i.e.* $1/3^{rd}$ of the Garman limit[38]) datasets to Xtrapol8. In the probed crystalline system, two monomers are found in the asymmetric unit (chains A and B) forming a functional dimer assembled by a four-helix bundle. Below, results and figures refer to monomer A; the results for monomer B are similar unless stated otherwise.

Reference (PDB entry 2VJA corresponding to AChE in complex with the substrate analogue collected with a dose of 3 MGy at 100) and triggered (PDB entries 2VJB corresponding to AChE in complex with the OTMA substrate analog collected with a dose of 9 MGy at 100) datasets were scaled anisotropically before the calculation of a $F_{obs}^{2VJB} - F_{obs}^{2VJA}$ (*i.e.*, $F_{obs}^{9MGy} - F_{obs}^{3MGy}$) *q*-weighted Fourier difference map and corresponding ESFAs for nine evenly-spaced occupancies between 0.1 and 0.9. Occupancies of the triggered states were determined using the *difference-map* method, and amounted to 0.2 (Supplementary Fig. 22). Again, we used the default truncate method to rescue negative ESFAs. As for the BAZ2BA-x538 test-case, carrying-out automatic refinement with the reference model was inappropriate since the reference and triggered states do not feature the same ligands in the active site gorge (substrate-analogue vs. radiolytic products). The $F_{obs}^{2VJB} - F_{obs}^{2VJA}$ *q*-weighted map calculated by Xtrapol8 is as featureful as that reported earlier, which was calculated in CNS using the original *q*-weighting script written by Thomas Ursby[9] (Supplementary Fig. 23).

Structure factor extrapolation allowed us to expand on the insights obtained in the initial study. Indeed, the $2mF_{extrapolated}$-$DF_{calc}$ and $mF_{extrapolated}$-$DF_{calc}$ maps reveal the presence of four radiolytic products in the active site: (i-ii) carbocholine and ethanal, from the radiation induced cleavage of the C5-C6 bond in the substrate analogue; and (iii-iv) two $CO_2$ molecules, from the decarboxylation of E199 and catalytic E327. Several residues in the active site are present in alternate conformations. Notably, two conformations are seen for carbocholine. In both, the choline moiety interacts by cation-π interactions with W84 and F330, yet each conformer coincides with different F330 conformations and different water networks. The first carbocholine conformer has its carbon chain pointing towards catalytic S200, while that of the second conformer points towards W432 and Y442, following a 180° rotation around the choline nitrogen position. Importantly, neither of the two carbocholine binding modes corresponds to that reported earlier for thiocholine [35] and choline,[36] either due to the change in their terminal atom (C, S, O, respectively) or to the void left by the decarboxylation of E199. Two close conformations are as well observed for the ethanal product, which is best modelled as either bound covalently to the catalytic S200 (conformer A, with the ethanal oxygen atom H-bonding (2.7 Å) to A201(N) in the oxyanion hole) or at Van der Waals distances from its side chain (conformer B, with the ethanal oxygen atom pointing towards S226(O)). In both cases, the methyl group faces the acyl-biding pocket, but only in conformer B are the catalytic S200 and H440 H-bonded, despite the latter adopting a conformation different from the resting state on the two alternate conformations. Indeed, a ~60° change in H440 side chain conformation is observed, presumably as a result from the untethering from E327, due to decarboxylation of this residue and repositioning of the $CO_2$ in the vicinity of S226(OH). The distribution of alternate conformations allows a global picture to emerge, whereby the carbocholine with its carbon chain pointing towards S200, the covalently bound ethanal, the gorge-closing F330 conformer and the $CO_2$ molecule emanating from E199 decarboxylation coexist in a first conformer (conformer A), whereas after $CO_2$ exit, the carbocholine reorients, the ethanal is released from S200, F330 assumes its gorge-opening conformation and a change in the conformation of S200 restores H-bonding to H440 (conformer B). The first set of alternate conformers can be proposed as representative of the structure just after cleavage of the substrate analogue, whereas the second set could correspond to the subsequent step. At the peripheral site, decarboxylation of a glutamate is also observed (E278), with the radiolytically-produced $CO_2$ molecule remaining trapped in a groove lined by the side chains of W279 and Y70, and the main chain carbonyl and side chain of I275. This causes W279 and Y70 to adapt a double conformation, which might be the reason for the absence of the substrate analogue at the peripheral site. Rather, a double conformation PEG molecule can be fitted, which adopts different conformations in the two monomers of the asymmetric unit.

As suggested by the $F_{obs}^{2VJB} - F_{obs}^{2VJA}$ map, radiation-induced decarboxylation is most evident at the functionally important E199, E327 and E278. Refinement based on extrapolated maps nonetheless indicates that this process occurs throughout the protein. Indeed, in monomer A (monomer B), only 6

(9) aspartate residues over 24 (24), and 6 (10) glutamate residues over 33 (34; residues 486-489 are undefined in chain A) preserve their carboxylic groups. No clear pattern emerges in terms of the predicted pKa (calculated using propka in pdb2pqr[39]) or burying into the structure of these radiation resistant acidic residues (Supplementary Fig. 22 e). Nonetheless, we note that none of the glutamic acid residues preserving their carboxylic group (16 in total) and only two aspartic acid residues preserving their carboxylic group (15 in total) are involved in Coulombic interactions, suggesting a relation between the resistance to radiation damage of acidic residues and the absence of Coulombic interactions involving their side chains. $CO_2$ molecules could be fitted at 10 (7) loci in monomer A (monomer B), but only three of these are buried in the protein moiety, nearby E199, E278 and E327. It must also be noted that all disulfide bridges (three per monomer: C67-C94, C254-C265, C402-C521) are ruptured in the extrapolated structures, as clearly pointed to by the $F_{obs}^{2VJB} - F_{obs}^{2VJA}$ Fourier difference map and in agreement with Ref[41] (Supplementary Fig. 23 b, c, e and f). Surprisingly, the C254-C265 disulfide break could not be unambiguously modeled as two individual cysteines as no density for a sulfur atom at position 254 could be observed. The elongated electron density at position 265 could suggest an asymmetric breaking mechanism at the C254(Cβ)-C254(Sγ) bond, *i.e.*, not at the C254(Sγ)- C265(Sγ) bond (Supplementary Fig. 23 b and e), and remains to be further investigated.

The Shoot-and-Trap example offers illustration that more information can be extracted from refinement against ESFAs than from the analysis of a Fourier difference map alone. From the latter, it was rightfully concluded that exposure to high X-ray doses leads to i) breakage of disulfide bridges; ii) decarboxylation of E199, E278 and catalytic E327; and (iii) cleavage of the non-hydrolysable substrate, with (iv) repositioning of the radio-produced choline adduct at 100 K. Upon structure refinement in ESFAs, we show that not only is the non-hydrolysable substrate analogue cleaved, but this cleavage occurs at the C5-C6 bond, resulting in the production of carbocholine and ethanal. The carbocholine reorients and adopts two as yet unobserved conformations in the active site, likely due to trapping of $CO_2$ at the bottom of the gorge, following decarboxylation of E199. One of these shows the carbon chain pointing towards the catalytic S200, and may represent the structure of the protein just after photocleavage of the substrate analogue. The other conformer has its carbon chain pointing towards Y442, following a 180° rotation around the choline position. The two conformers coincide with different conformations of F330 and different water networks. Two close conformations are as well observed for the ethanal product, which is either bound covalently to the catalytic S200 (conformer A) or found at Van der Waals distances from its side chain (conformer B).

#### Supplementary Discussion

Here we reported the design and use of Xtrapol8, a program aimed at facilitating the determination of low-occupancy state structures in time-resolved, kinetic and ligand-binding crystallography. Examples presented above demonstrate that Xtrapol8 can be useful in a wide variety of cases, from ligand-binding to kinetic and time-resolved crystallography.

Xtrapol8 was designed with view to address three main issues. Firstly, we wanted to make available to a wide community of crystallographers a tool enabling eased calculation of Bayesian-statistics weighted Fourier difference maps and ESFAs, and this independent of their crystallographic expertise. To this end, our software had to be user-friendly and highly-automated, requiring a minimal input from inexperienced users while allowing expert crystallographers to tweak the calculations with their preferences. Hence, many parameters such as resolution cutoffs, scaling parameters, weighting schemes, as well as parameters for difference map exploration or refinement can be defaulted or adjusted. To further ease its usage, Xtrapol8 can be controlled through an intuitive graphical user interface as a frontend to the command line version, which we hope will reduce the barriers for inexperienced crystallographers to use extrapolation methods. Secondly, as different approaches for the calculation of Fourier difference map and extrapolation of structure factor amplitudes have been proposed, we sought to make all of them available within a single tool, so as to allow users to compare results obtained using different strategies. Indeed, we observed that depending on the case, a specific strategy might be more successful than another; Xtrapol8 allows testing all methods within a single run. Thirdly, we aspired to address the issue of negative ESFAs, whose percentage increases as the occupancy of the intermediate state decreases, degrading the quality of extrapolated maps while making reciprocal-space and real-space refinement unstable. Presence of negative ESFAs can be unnoticed when using custom-written scripts, leaving their treatment – most often their elimination – to the refinement software that is used downstream. As shown by our examples, the best approach depends on the actual data, the total number of negatives and if any bias can be afforded. If the negative reflections are simply rejected, the effective completeness of the data used in refinement can strongly decrease, leading to a degradation in map quality and refinement results. Last, with view to increase overall reproducibility and transparency, we wanted to provide methods to objectively estimate the occupancy of the triggered state(s) based exclusively on the diffraction data. Two such methods are accordingly available within Xtrapol8. The *difference-map* method maximizes peak height in the extrapolated difference maps, whereas the *distance-analysis* method exploits changes in atomic positions in the triggered state models produced by reciprocal- and/or real-space refinement against ESFAs and/or extrapolated electron density maps, respectively, with respect to the reference model. Ideally, both occupancy estimation methods should be used to enable cross-validation of the occupancy estimation. However, usage of the latter method requires proper refinement settings and correct modelling of the extrapolated density map. In some cases, illustrated by our three examples, the automatic reciprocal-space and real-space refinement should not / cannot be carried out before manual intervention on the model, hence only the *difference-map* method will initially be usable. Nevertheless, a script is provided to re-run the *distance-analysis* method in stand-alone mode so that the user may intervene manually to produce a first model of the triggered state with which the refinements can be relaunched and occupancy determined via the *distance-analysis* method. Importantly, the *distance-analysis* method may be the only valid option in cases where atomic motions are small, data are weak, or multiples excited states coexists, leading to a flattening of peaks in the Fourier difference map.

Of important note, Xtrapol8 can also be used for the evaluation of ligand binding and fragment screening assays, complementing already existing pipelines such as PanDDA.[29] The latter decides on the presence of a potential ligand based on the electron density distribution and then uses an ensemble of ligand-free electron density maps to locate the ligand through real-space electron density deconvolution. In contrast, Xtrapol8 uses a single *a priori* ligand-free data set and bases all occupancy-estimations and structure-refinements on SFA differences. The two methods are thus highly complementary. Indeed, the PanDDA strategy is less dependent on crystallographic isomorphism and generates nearly noise-free local electron density maps, but the calculation does not allow fast feedback and requires an ensemble-based refinement in the initial untreated data. Contrastingly, Xtrapol8 allows to calculate a Fourier difference map in a matter of minutes, allowing real-time feedback, and the ESFAs can be used to refine the structure using standard protocols.

Our examples illustrate how important occupancy estimation can be for the interpretation of structural results. In the case of crystallography experiments where the completeness and multiplicity of the data is high (*e.g.* serial crystallography), jackknife resampling could be implemented[40] offering a means to increase the statistical significance of the occupancy estimate. To do so, multiple random (reference and triggered) datasets would have to be produced, each lacking a given fraction of the collected frames, enabling to run Xtrapol8 multiple times on slightly different data, and to base occupancy determination on the distribution of the estimated occupancy for each pair of jackknifed datasets. While jackknife resampling is not part of the current Xtrapol8 release, it will be incorporated soon.

#### Supplementary Figures and Tables

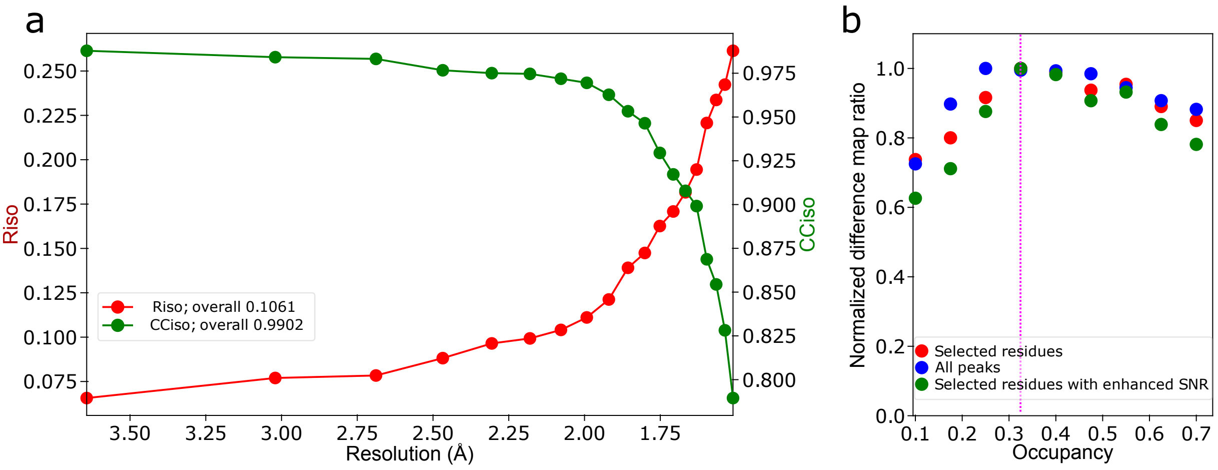


**Supplementary Fig. 1 | a,** mEos4b isomorphism, expressed as Riso and CCiso, in function of resolution. **b,** mEosb4b occupancy determination after a first Xtrapol8 run in '*fast-and-furious'* mode. The automatically annotated occupancy value is indicated by a magenta dashed line.

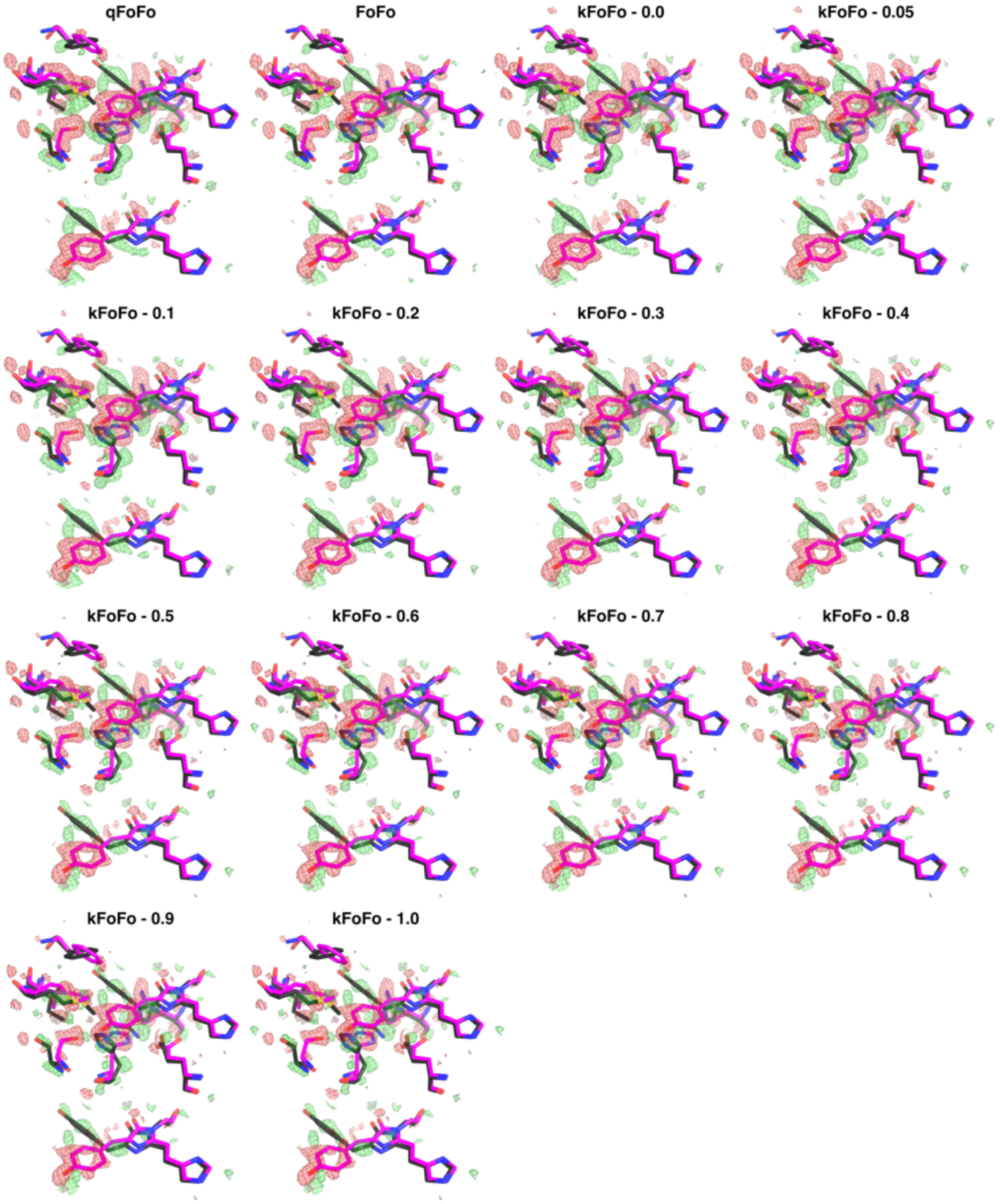


**Supplementary Fig. 2 |** mEos4b chromophore environment (top) and chromophore only (bottom) in the *red-on* and *red-off* state (magenta and gray sticks, respectively) superposed with Fourier difference maps (contoured at ± 3 σ) calculated with the *q*-weighting, no-weighting and *k*-weighting schemes. For *k*-weighting, the additional factor for outlier downweighing is also indicated.

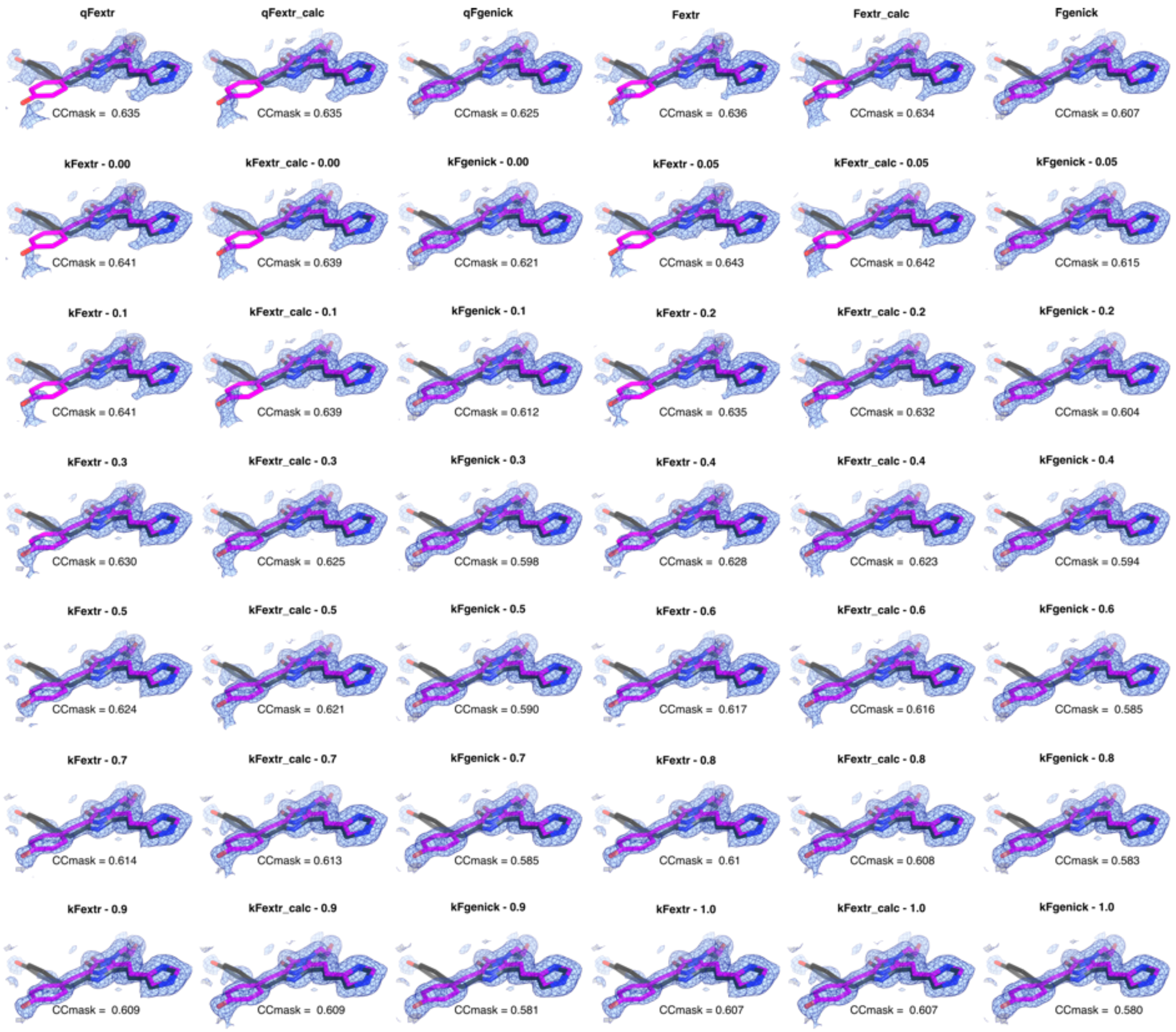


**Supplementary Fig. 3 |** mEos4b chromophore in the *red-on* and *red-off* state (magenta and gray sticks, respectively) superposed on the $2mF_{extrapolated}$-$DF_{calc}$ electron density map calculated for an occupancy of 0.325, contoured at 1 σ, for each type of ESFA. In case of *k*-weighting, also the additional scale factor is indicated. For each map, the CCmask is indicated which is calculated using the chromophore and surrounding residues.

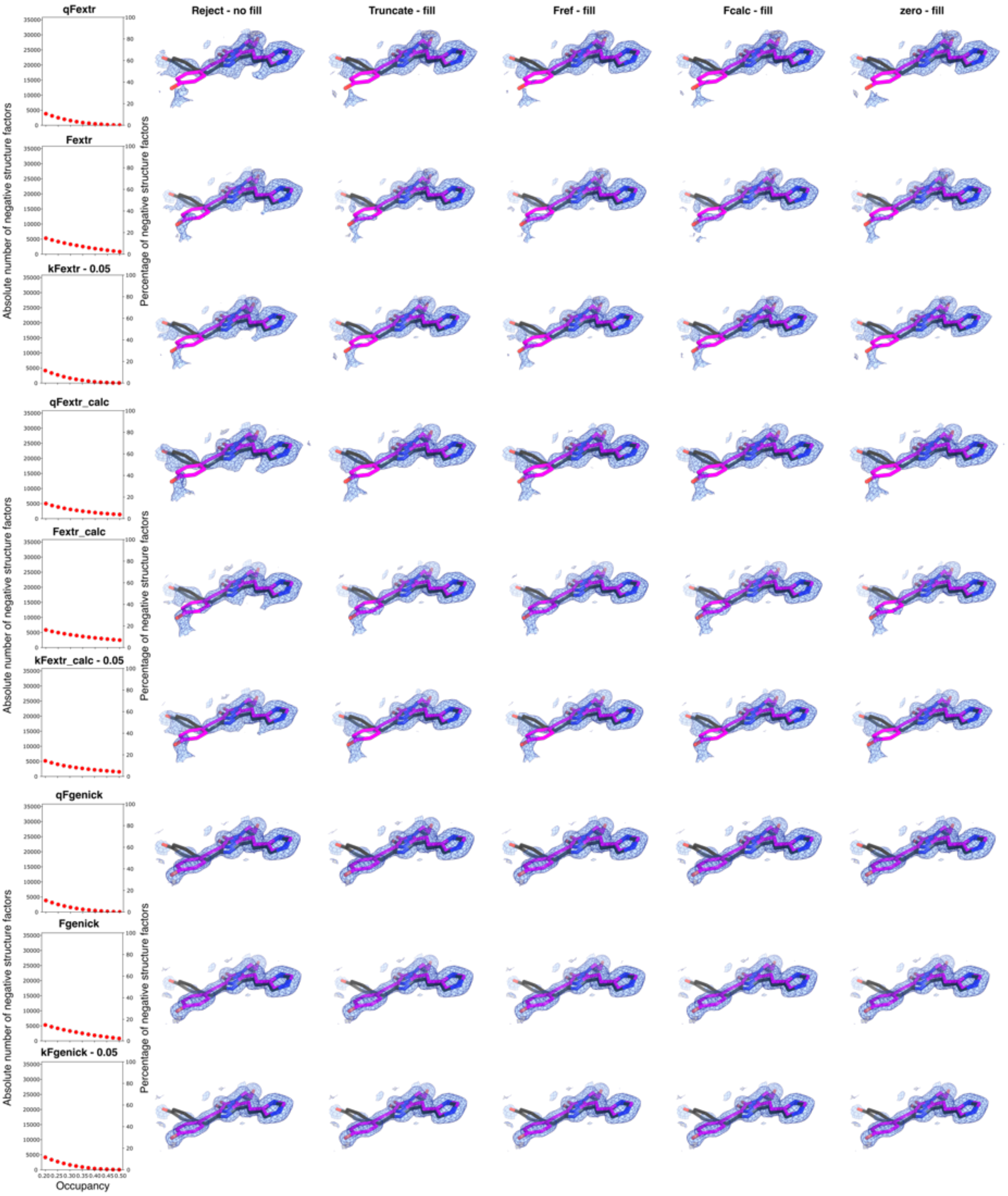


**Supplementary Fig. 4 |** Number of negative ESFAs (first column) and $2mF_{extrapolated}$-$DF_{calc}$ electron density map calculated for an occupancy of 0.325 using different ways to treat negative ESFAs (columns) and ESFA calculation schemes (rows), contoured at 1 σ for the mEos4b chromophore. For *k*-weighting an outlier scale factor of 0.05 was used.

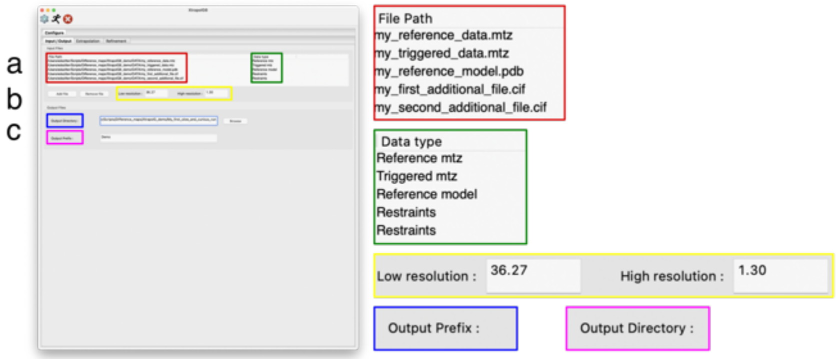


**Supplementary Fig. 5 |** Panel of the Xtrapol8 graphical user interface for input and output management, with zoom-in of several fields (colored boxes). **a,** input files and specification of their type (reference mtz, triggered mtz, reference model and additional cif files if required). **b,** resolution limits can optionally be adapted. **c,** fields to alter the output directory and prefix for output files.

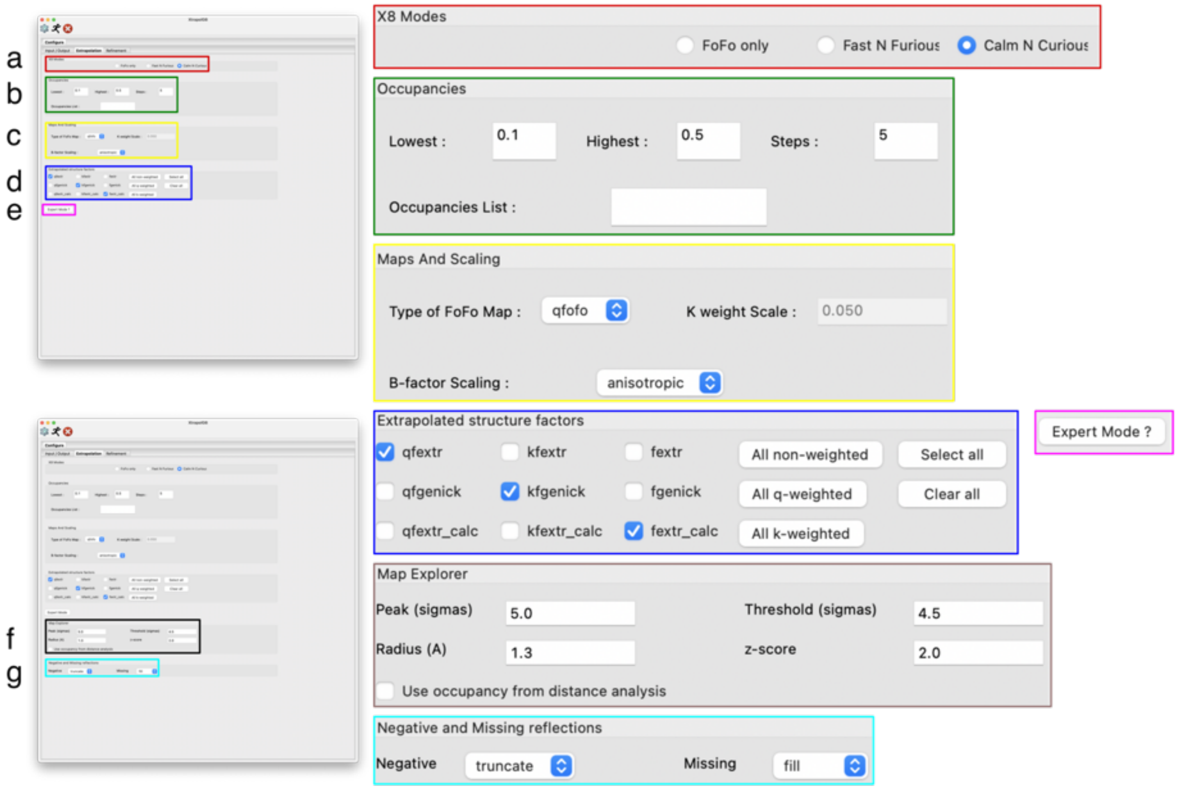


**Supplementary Fig. 6 |** Panel of the Xtrapol8 GUI to specify parameters and strategies for Fourier difference map and ESFAs calculation, with zoom-in of several fields (colored boxes). **a,** users can choose between 3 modes to run Xtrapol8: '*FoFo only*', '*fast and furious*' and '*calm and curious'*. Selection of a specific mode will automatically enable/disable certain parameters to alter. **b,** fields to alter the occupancies to be tested. This can be specified via a minimum and maximum value and number of steps in between or by a list with the values. **c,** in the "Maps and Scaling" fields, the user can precise if the Fourier map should be *q*-weighted, *k*-weighted or non-weighted, as well as how the reference and triggered datasets should be scaled. **d,** field to select the ESFA types. Multiple can be selected in a single run. **e,** more advanced options are offered to more experienced user and concern the parameter for analysis of the difference maps and occupancy estimation **(f)** and the treatment of negative and missing ESFAs **(g)**.

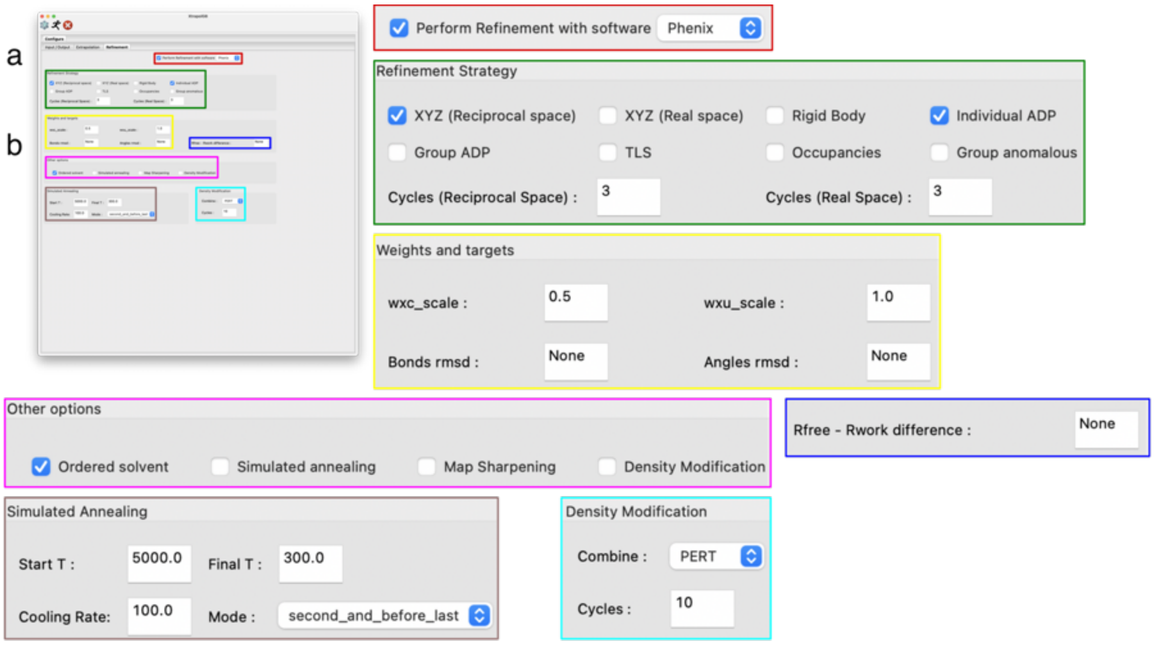


**Supplementary Fig. 7 |** Panel of the Xtrapol8 GUI concerning refinement with Phenix, with zoom-in of several fields (colored boxes). **a,** when the "Perform refinement with" checkbox is ticked, reciprocal and real-space refinement can be carried out using phenix.refine and phenix.real_space_refine. **b,** several options for reciprocal-space refinement with phenix.refine can be altered. The number of refinement cycles in phenix.real_space_refine can also be set.

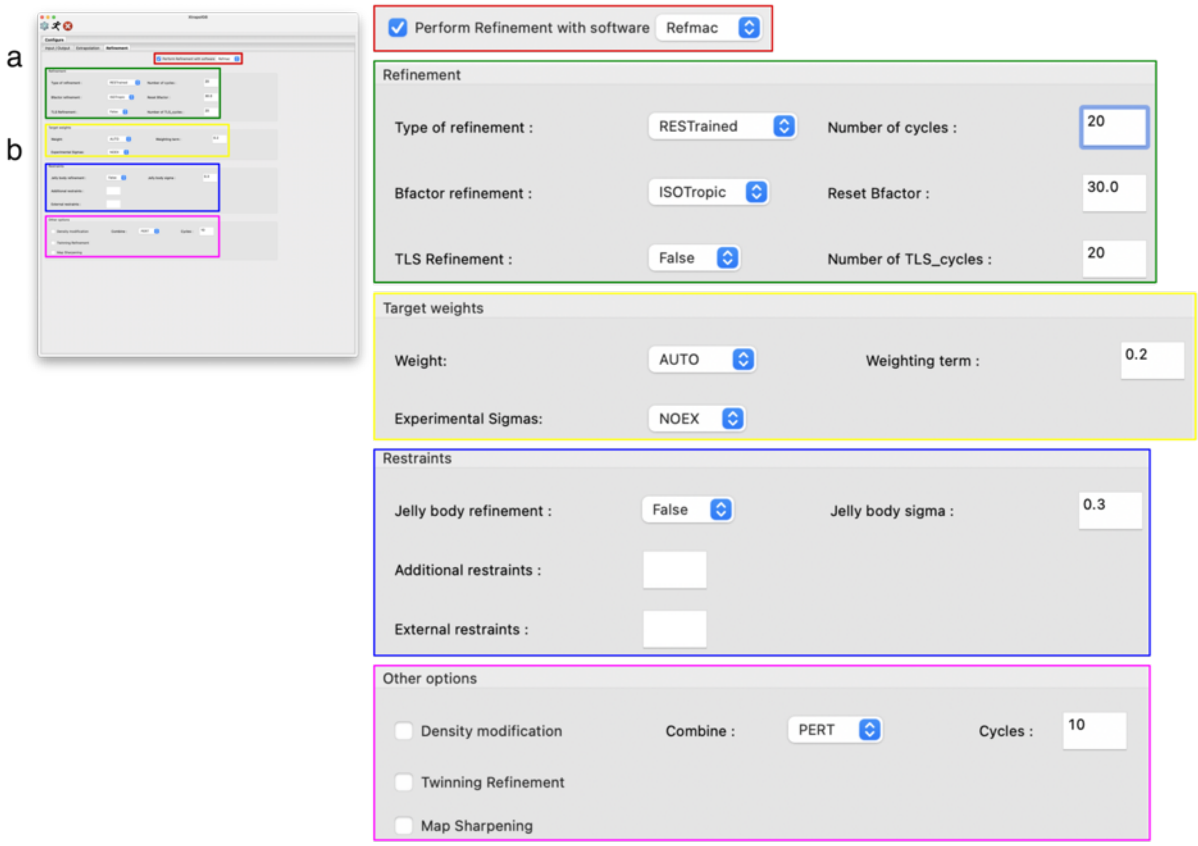


**Supplementary Fig. 8 |** Alternative panel of the Xtrapol8 GUI concerning refinement with CCP4, with zoom-in of several fields (colored boxes). **a,** when the "Perform refinement with" checkbox is ticked, reciprocal and real-space refinement can be carried out using Refmac and Coot. **b,** several options for reciprocal-space refinement with Refmac can be altered.

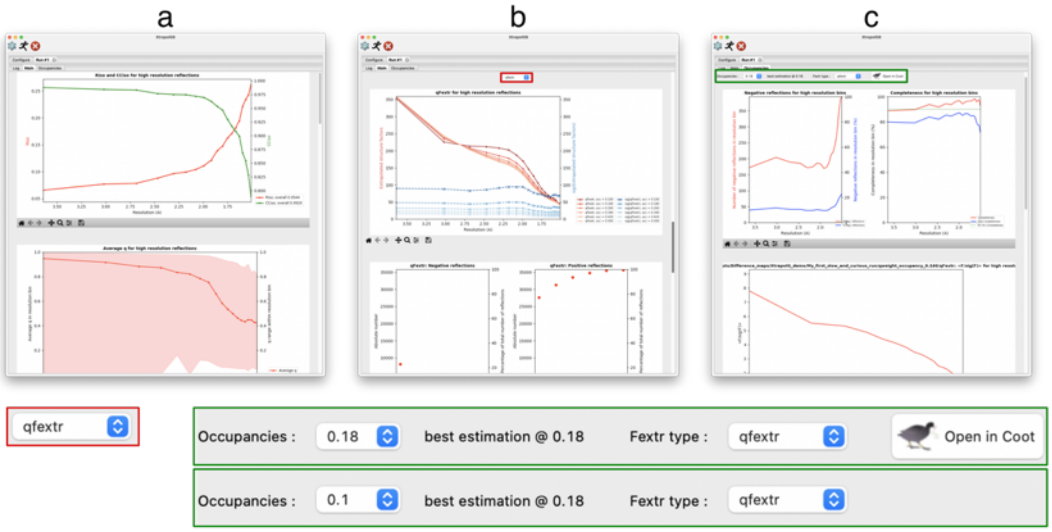


**Supplementary Fig. 9 |** The results of an Xtrapol8 run using the GUI, with zoom-in of specific fields (colored boxes). A first tab contains a print of the log-file (not shown). Two other tabs contain the main output or the output considering the occupancies. **a-b,** snippets of the main output tab. **a,** the top part contains general figures, such as the one indicating the isomorphism or the value of the *q*-weights as a function of the resolution. **b,** the bottom part contains information of ESFAs for which the type should be selected via the dropdown menu (red), *e.g.* signal strength and number of negative and positive ESFAs. **c,** the occupancies tab contains information concerning the ESFAs at each of the tested occupancies, which both have to be selected by a dropdown menu (green). The best estimate of the occupancy for the ESFA type will be indicated and models and maps for that occupancy can be automatically opened in Coot.

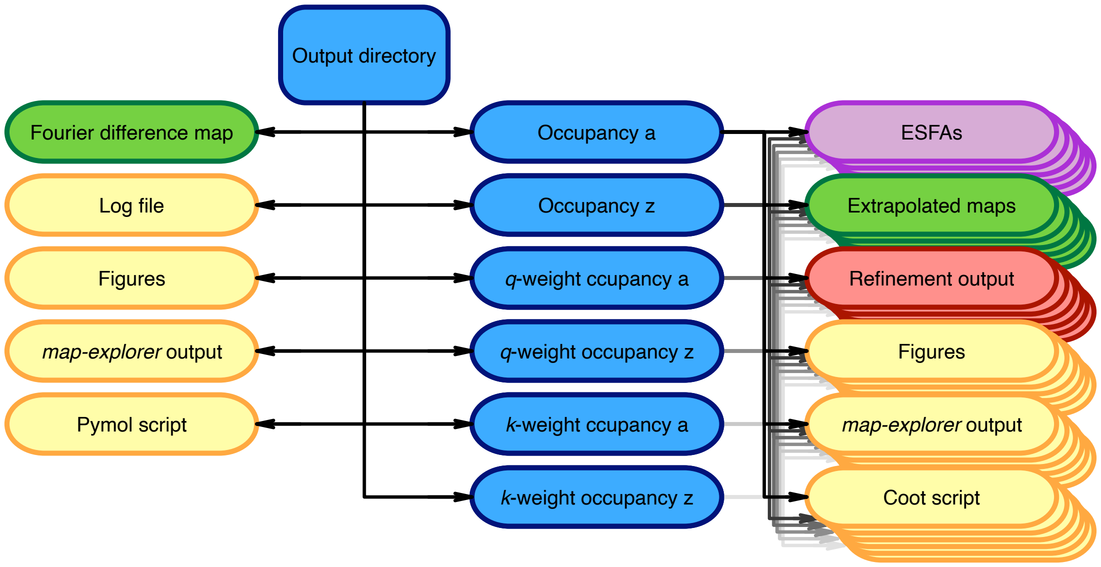


**Supplementary Fig. 10 |** Schematic representation of the Xtrapol8 output directory. (Sub)directories are shown in blue, electron density maps in green, structure factors in purple, refinement output in red and all other files intended to aid in interpretation and decision making in yellow. Subdirectories are created for each tested occupancy a to z, with the directory name specifying whether Bayesian weighting (*q*/*k*-weighting) or no weighting was used in the calculation of ESFAs.

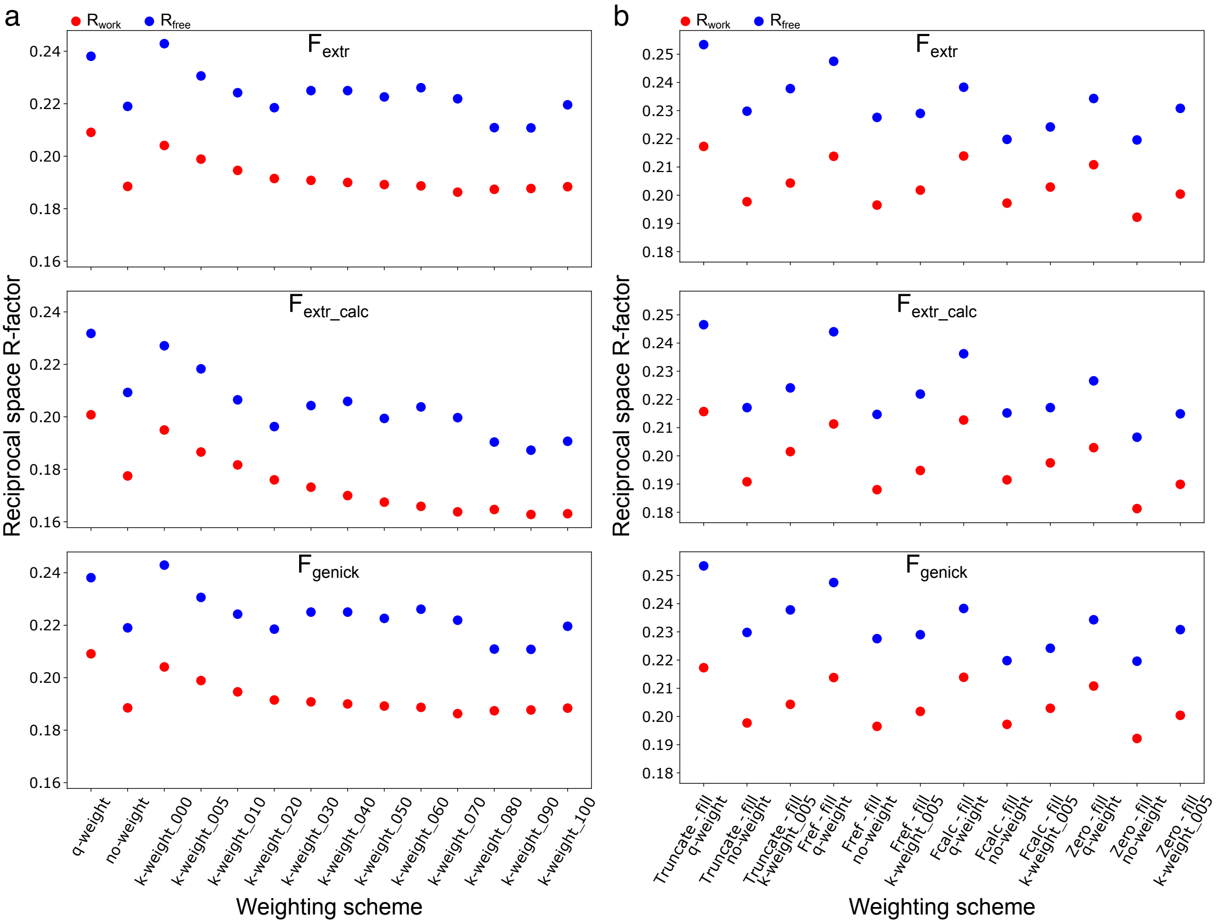


**Supplementary Fig. 11** | $R_{work}$ (red) and $R_{free}$ (blue) values after reciprocal-space refinement of the published red model (PDB entry 6GP1) in various types of ESFAs and different weighting schemes using the *refiner.py* script. **a,** negative ESFAs were rejected and R-factors are reported for *q*-weighting, no-weighting and *k*-weighting with various *k*-weighting outlier scale factors. **b,** the truncate-, $F_{obs}^{reference}$-, $F_{calc}^{reference}$- and zero-methods are used to estimate negative ESFAs that are *q*-weighted, non-weighted or *k*-weighted (with outlier scale factor of 0.05).

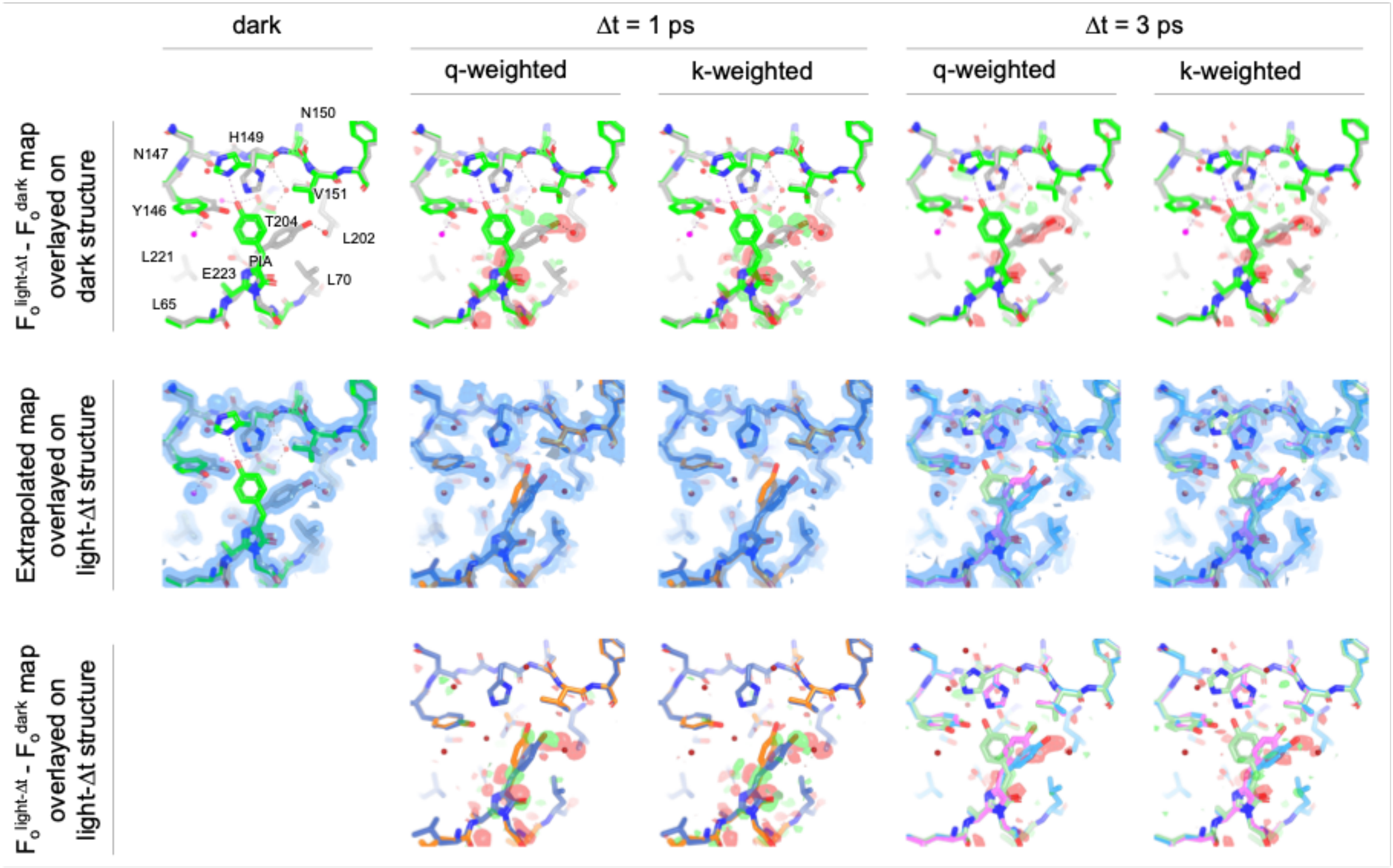


**Supplementary Fig. 12 |** Summary of Xtrapol8 outcome on the extraction of ps lived intermediates states in rsEGFP2. A dataset collected on crystalline rsEGFP2 in the OFF state was used as the reference dataset (laser-OFF), while two datasets collected at 1 and 3 ps post-pumping by a UV laser fs-pulse served as the triggered datasets (laser-ON). The top row shows the laser-OFF state structure, which features 90% of molecules in the fluorescent OFF-state (grey) and the remainder in the fluorescent ON-state (green), with the *q*/*k*-weighted Fourier difference maps overlaid. The middle row shows the $2mF_{obs}$-$DF_{calc}$ (laser-OFF) and *q*/*k*-weighted $2mF_{extrapolated}$-$DF_{calc}$ (laser-ON) electron density maps overlayed on the refined laser-ON models (conformers are depicted with different colors). The bottom row shows a superposition of the laser-ON models superposed on the *q*/*k*-weighted Fourier difference maps.

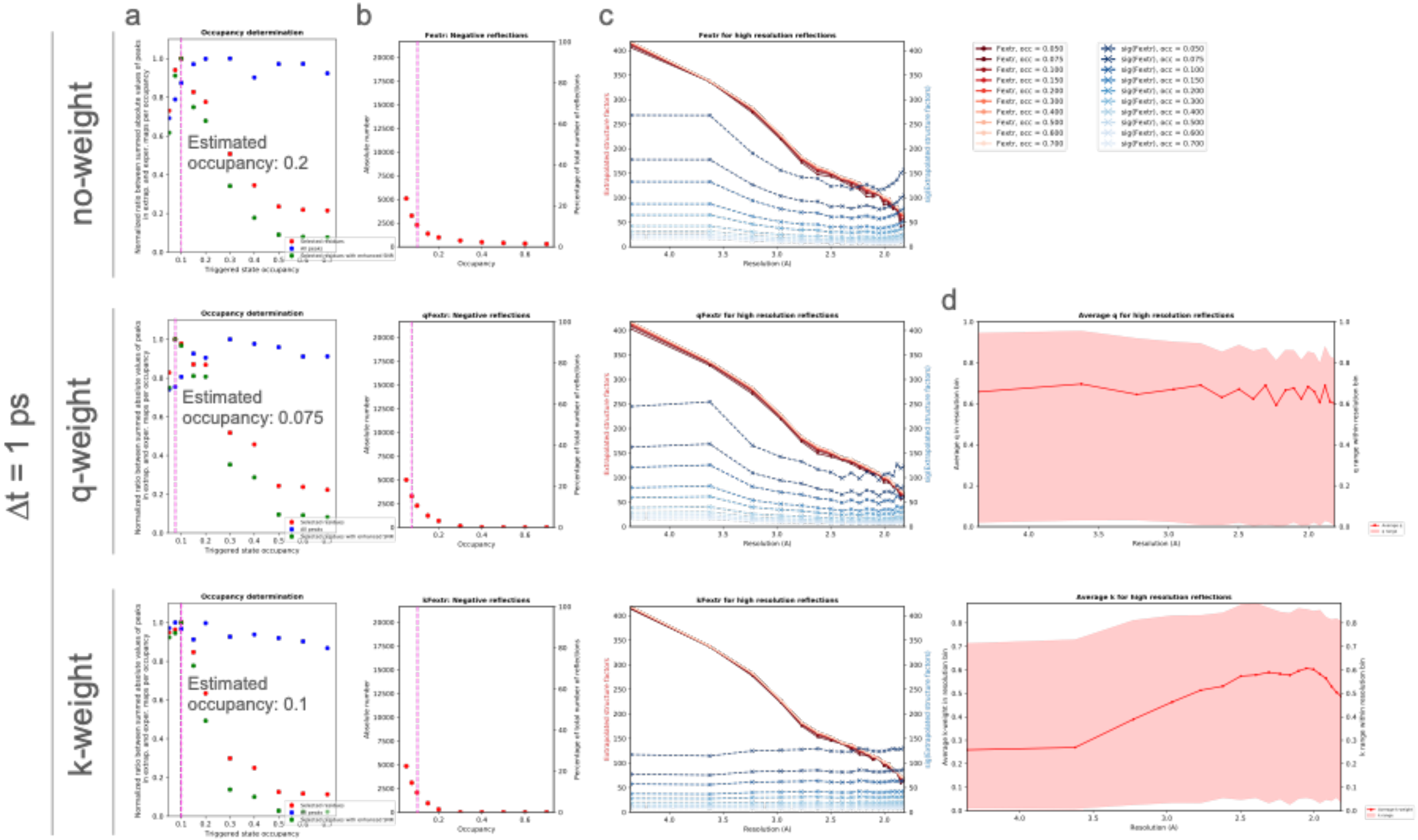


**Supplementary Fig. 13 |** Effect of Bayesian-statistics weighting of ESFA computed for various occupancies from the reference and Δt=1 ps data. The top, middle and bottom rows show results with no weighting, *q*-weighting or *k*-weighting of EFSAs. **a**, Bayesian weighting of ESFAs hardly influences the outcome of occupancy determinations carried out using the difference map method. **b**, Bayesian weighting of ESFAs only slightly reduces the amount of negative reflections. In (**a**) and (**b**), the occupancies determined by the difference map method are highlighted by a pink shaded line. **c**, Plots of mean ESFA and corresponding σ(ESFA) as a function of resolution. **d**, Plots of *q*-weights and *k*-weights average value and range of values as a function of resolution.

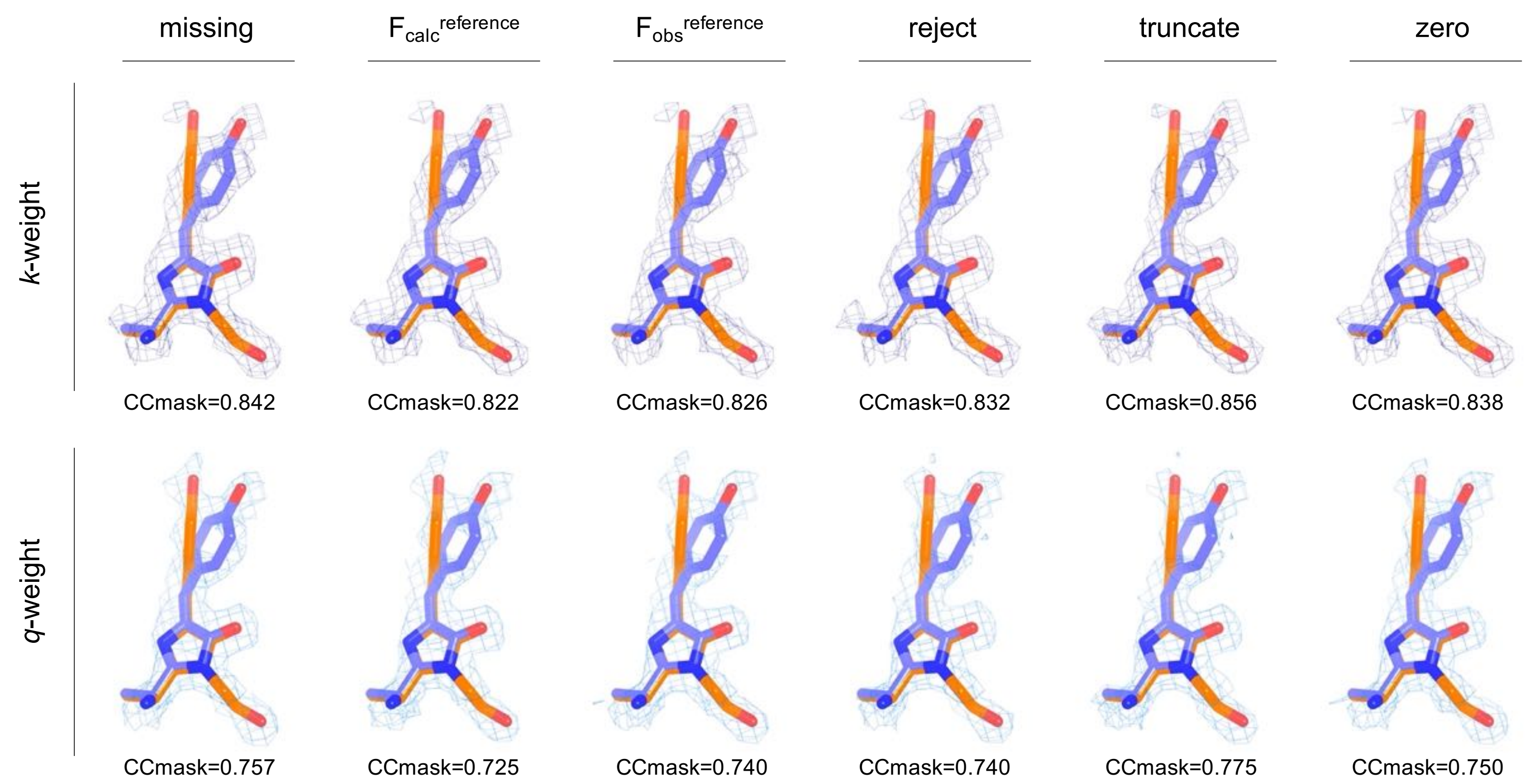


**Supplementary Fig. 14 |** Illustration of the effect of negative-ESFA handling on the extrapolated maps calculated for the Δt = 1ps time delay. The initial $2mF_{extrapolated}$-$DF_{calc}$ maps obtained using the various negative-ESFAs handling-approaches implemented in Xtrapol8 and a triggered-state occupancy of 7.5 % are overlaid on the laser-ON model refined on the basis of the *k*-weighted ESFA produced with the truncate method. For clarity, only the chromophore and the corresponding extrapolated electron density are shown. The overall CCmask of the model is indicated, which was calculated using all atoms in the models. The CCmask for the *q*-weighted extrapolated maps and models is lower than that for the *k*-weighted counterparts, possibly due to damping of structural differences due by the *k*-weighting procedure, as compared to *q*-weighting. Regardless, the CCmask is higher for the extrapolated maps calculated from ESFAs produced with the truncate method, showcasing the importance of rescuing negative ESFAs.

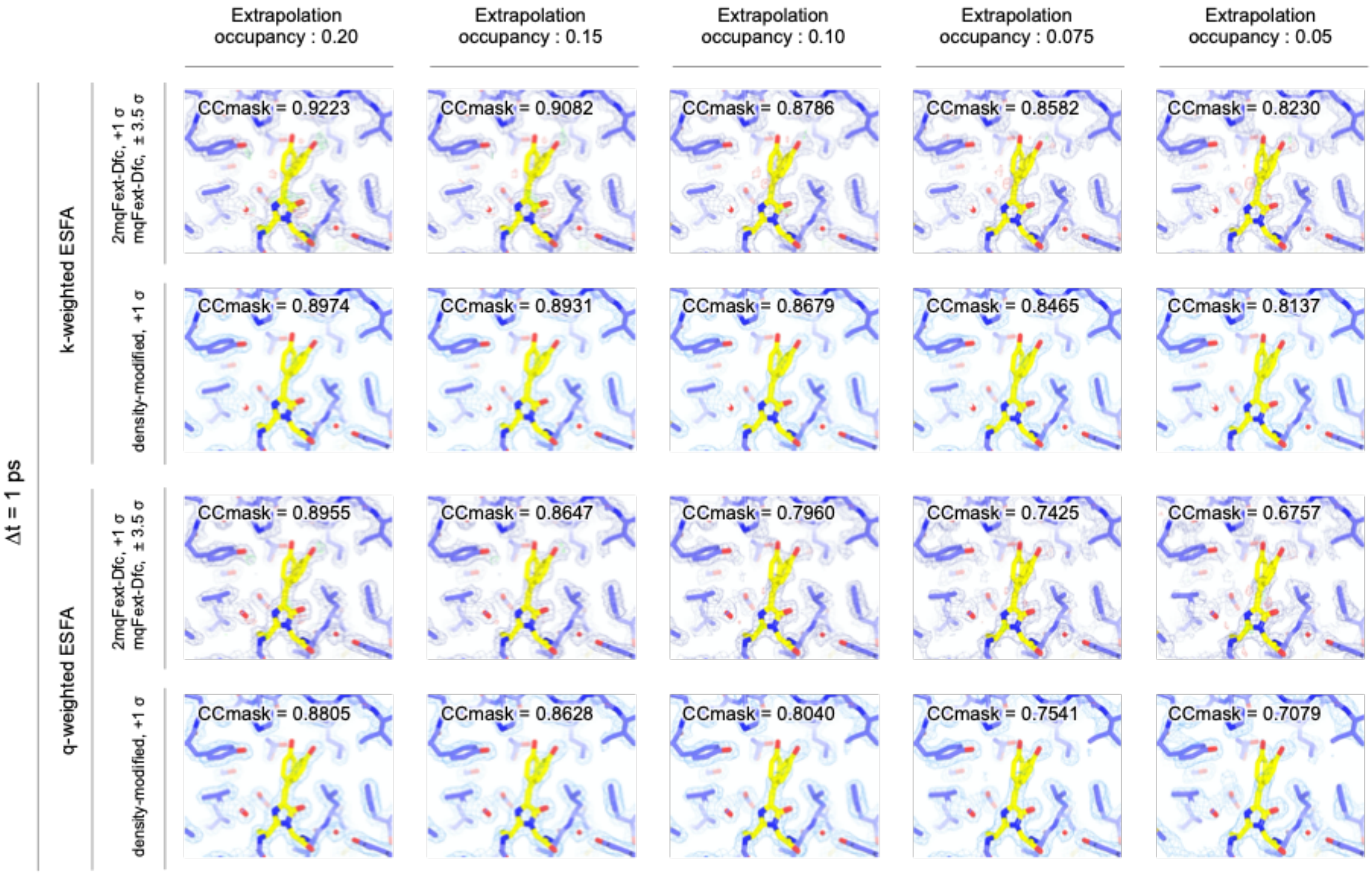


**Supplementary Fig. 15 |** Evolution of features and noise in the *k*-weighted and *q*-weighted $2mF_{extrapolated}$-$DF_{calc}$ initial and density modified maps computed at various extrapolation occupancies from the laser-OFF and laser-ON Δt=1 ps datasets. While density modification slightly benefits map quality in the case of *q*-weighted ESFAs, it slightly decreases it in the case of *k*-weighted ESFAs. Thus, while it sounds reasonable that density modification should improve the quality of the maps, it is not always the case in practice and hence the opportunity of basing real-space refinement on these should be verified on a case-by-case basis.

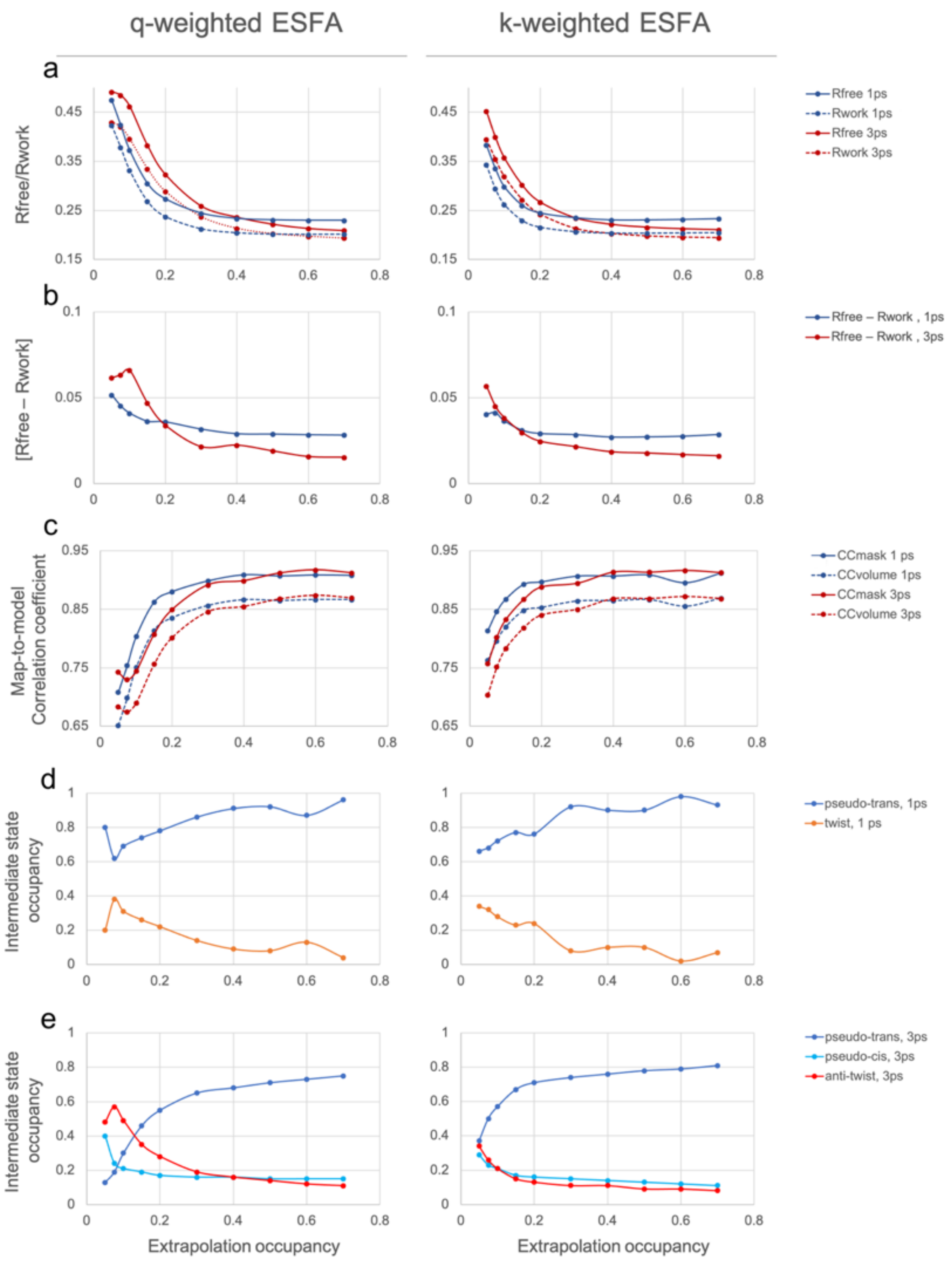


**Supplementary Fig. 16 |** Automatic refinement of the laser-ON models of rsEGFP2 using the *refiner.py* script of Xtrapol8. For both the Δt=1 ps and Δt=3 ps laser-ON models, automatic refinement was unable to account for the large change in chromophore conformation, hence, manual intervention was required. For each time delay, a conservative model was produced on the basis of features in the *k*-weighted $2mF_{extrapolated}$-$DF_{calc}$ map computed for an occupancy of 0.2, which was then subjected to refinement against *q*-weighted (left column) and *k*-weighted ESFAs (right column) calculated for all occupancies. **a,** $R_{work}$ and $R_{free}$ values of the extrapolated structures after 10 cycles of reciprocal-space refinement including positional and occupancy refinements; results are shown for the Δt=1 ps (blue) and the Δt=3 ps (red) datasets. **b,** $R_{work}$ - $R_{free}$ difference values; to ensure that this value remains in the 3-5% range even at lower occupancies, the wxc_scale value was set to 0.0000002. **c,** CCmask and CCvolume values after real-space refinement in the reciprocal-space refined extrapolated density maps. **d-e,** Refined occupancies determined for the two conformers upon refinement against *q*-weighted and *k*-weighted ESFAs for the Δt=1 ps **(d)** and Δt=3 ps **(e)** datasets.

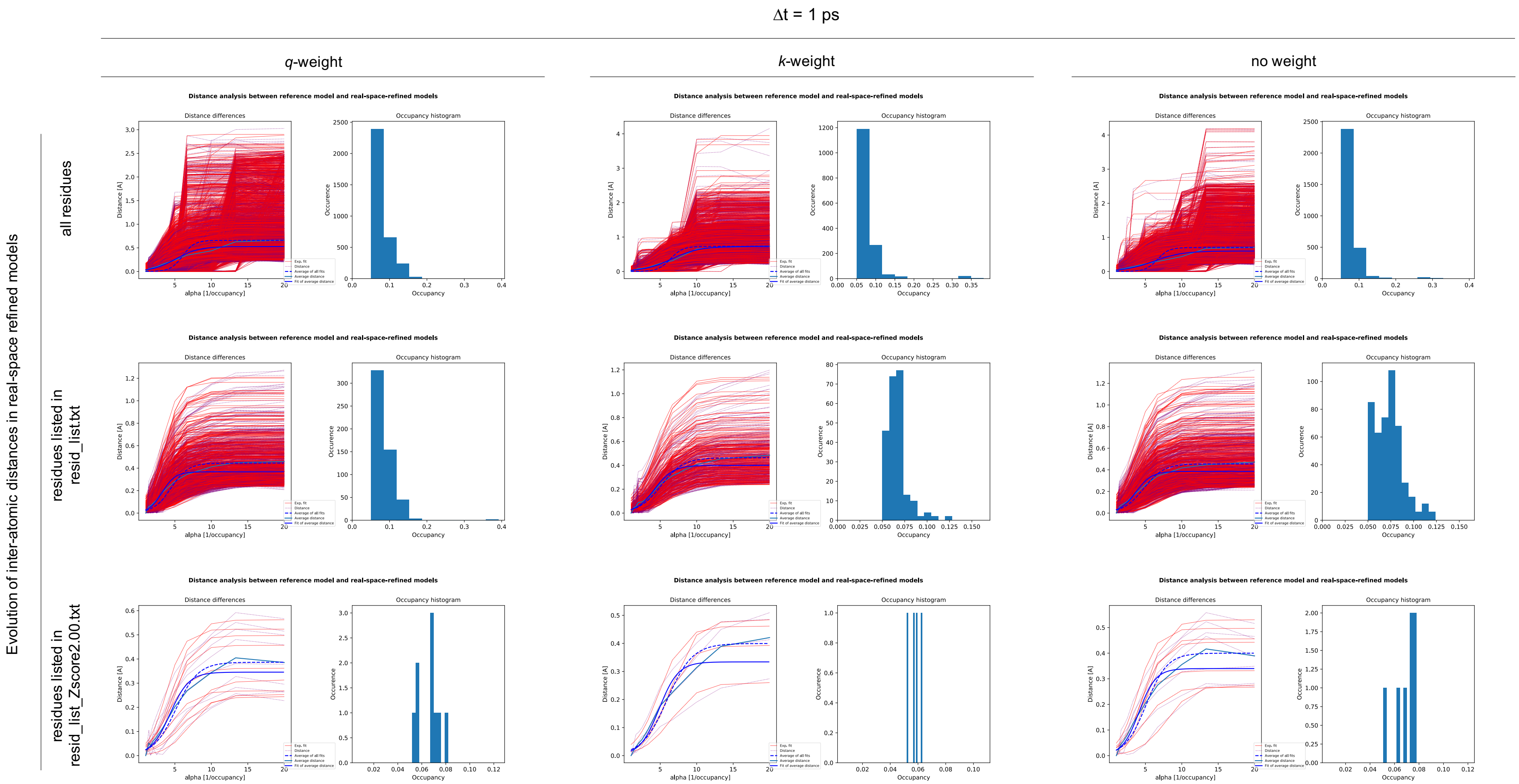


**Supplementary Fig. 17 |** Occupancy determination for the Δt = 1 ps dataset based on the distance analysis method. The distance analysis method estimates the correct occupancy by fitting with a sigmoid the evolution of interatomic distances as a function of all tested occupancies and then retrieving the occupancy value for which 99% of the plateau is reached.

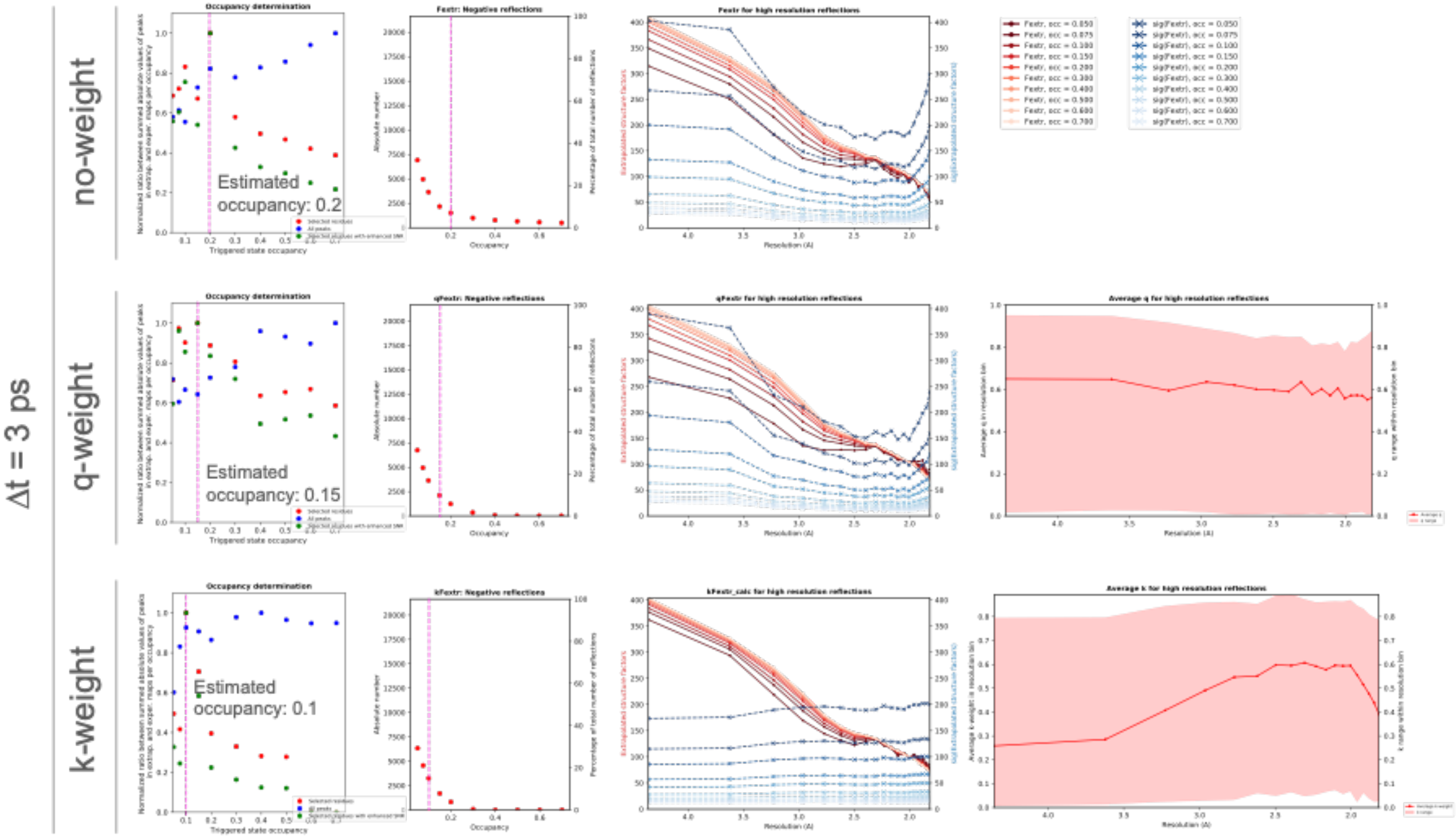


**Supplementary Fig. 18 |** Effect of Bayesian-statistics weighting of ESFA computed for various occupancies from the reference and Δt=3 ps data. The top, middle and bottom rows show results with no weighting, *q*-weighting or *k*-weighting of EFSAs. First column: Bayesian weighting of ESFAs hardly influences the outcome of occupancy determinations carried out using the difference map method. Second column: Bayesian weighting of ESFAs only slightly reduces the amount of negative reflections. In the first and second column, the occupancies determined by the difference map method are highlighted by a pink shaded line. Third column: Plots of mean ESFA and corresponding σ(ESFA) as a function of resolution. Last column: Plots of *q*-weights and *k*-weights average value and range of values as a function of resolution.

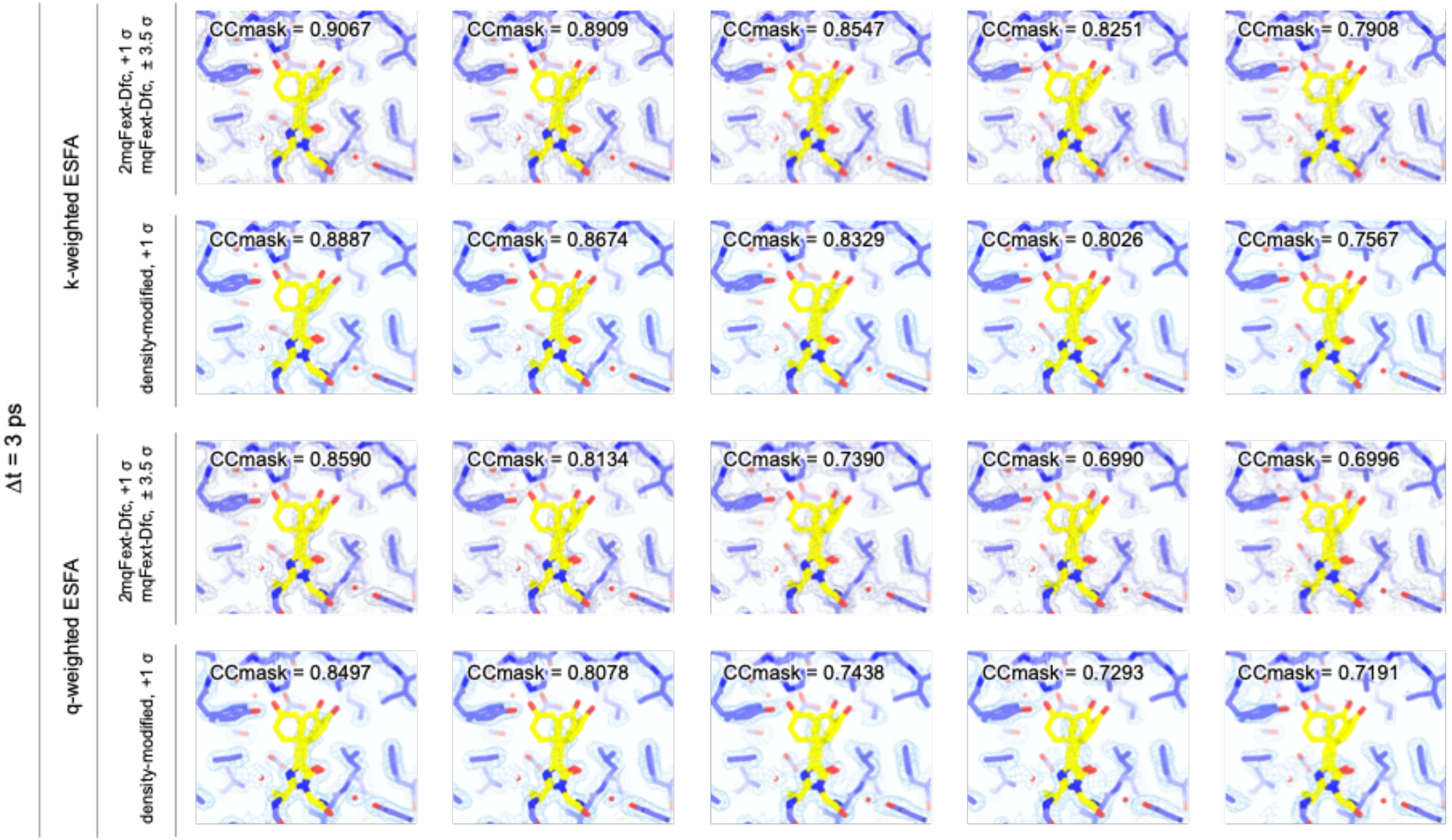


**Supplementary Fig. 19 |** Evolution of features and noise in the *k*-weighted and *q*-weighted $2mF_{extrapolated}$-$DF_{calc}$ initial and density modified maps computed at various extrapolation occupancies from the laser-OFF and laser-ON Δt=3 ps datasets. While density modification slightly benefits map quality in the case of *q*-weighted ESFAs, it slightly decreases it in the case of *k*-weighted ESFAs. Thus, while it sounds reasonable that density modification should improve the quality of the maps, it is not always the case in practice and hence the opportunity of basing real-space refinement on these should be verified on a case-by-case basis.

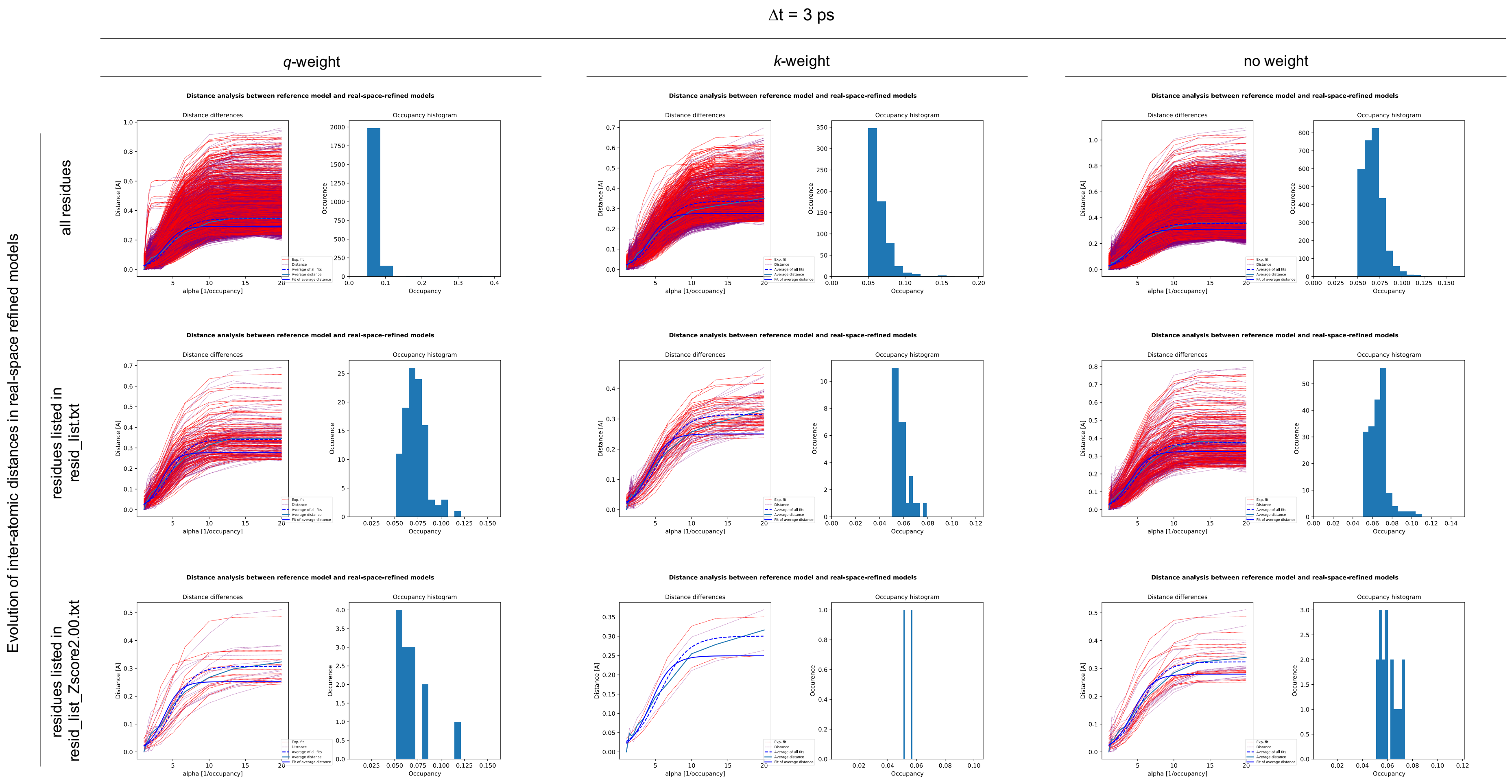


**Supplementary Fig. 20 |** Occupancy determination for the Δt = 3 ps dataset based on the *distance-analysis* method. This method estimates the correct occupancy by fitting with a sigmoid the evolution of interatomic distances as a function of all tested occupancies and then retrieving the occupancy value for which 99% of the plateau is reached.

.

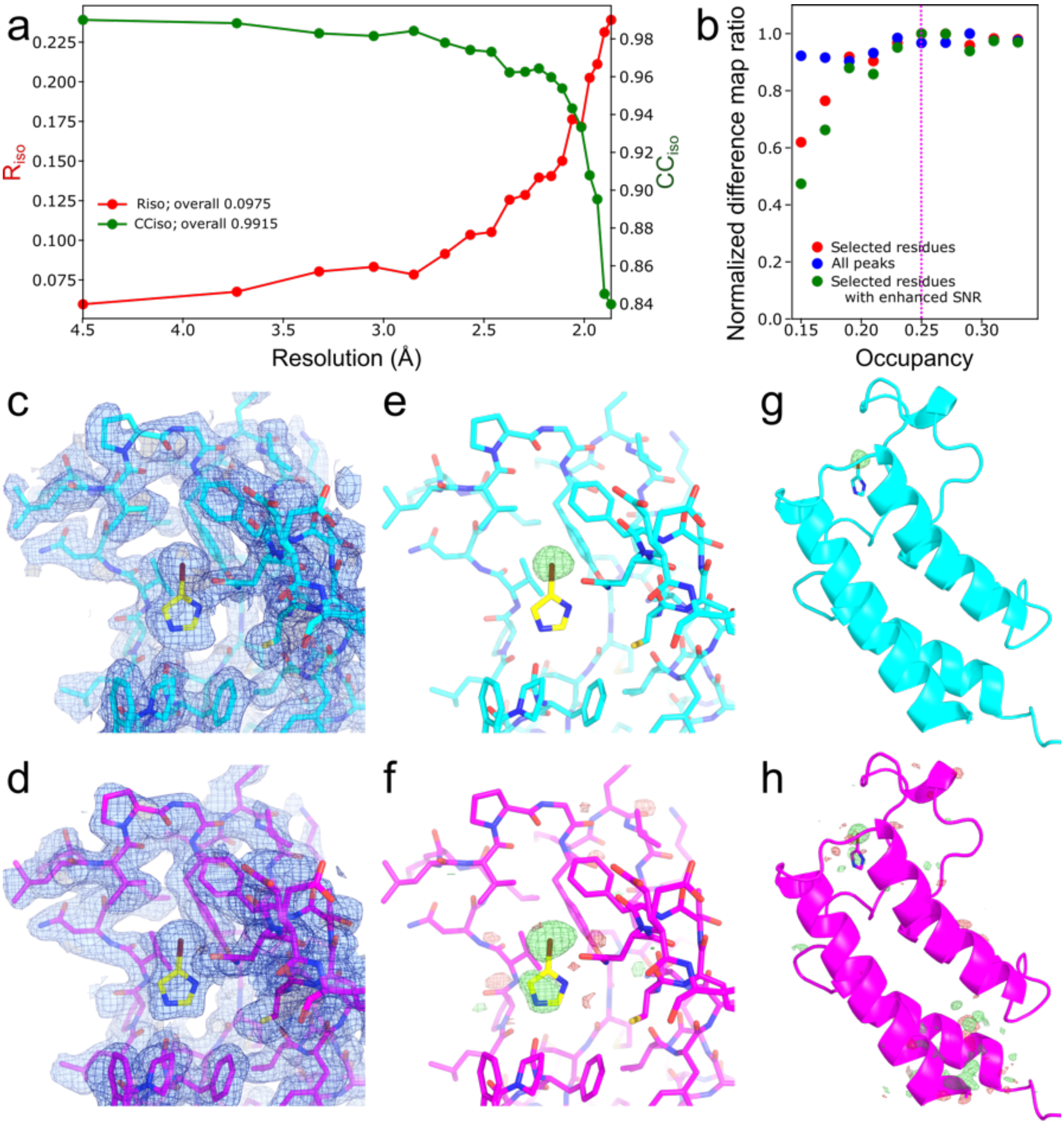


**Supplementary Fig. 21 |** Comparison of Xtrapol8 with PanDDA on BAZ2BA-x538. **a,** Isomorphism (Riso and CCiso) between the BAZ2BA-x538 and BAZ2BA-x645 data sets up to a high resolution of 1.8 Å. **b,** occupancy with the difference map method for the ligand using structure factor extrapolation (type: *q*Fextr). **c,** PanDDA event map for the ligand and surroundings, contoured at 1 σ. **d,** *q*-weighted $2mF_{extrapolated}-DF_{calc}$ electron density map with an occupancy of 0.25 for the ligand and environment, contoured at 1 σ. **e,** PanDDA z-map for the ligand, contoured at ± 4 σ. **f,** *q*-weighted Fourier difference map for the ligand, contoured at ± 4 σ. **g,** PanDDA z-map for the complete protein, contoured at ± 4 σ. **h,** *q*-weighted Fourier difference map for the complete protein, contoured at ± 4 σ. The model in cyan and magenta are the BAZ2BA-x645 model wherein the 1,2-ethanediol is replaced by the 4-bromoimidazole ligand (yellow) based on the PanDDA event or on extrapolated electron density maps, respectively.

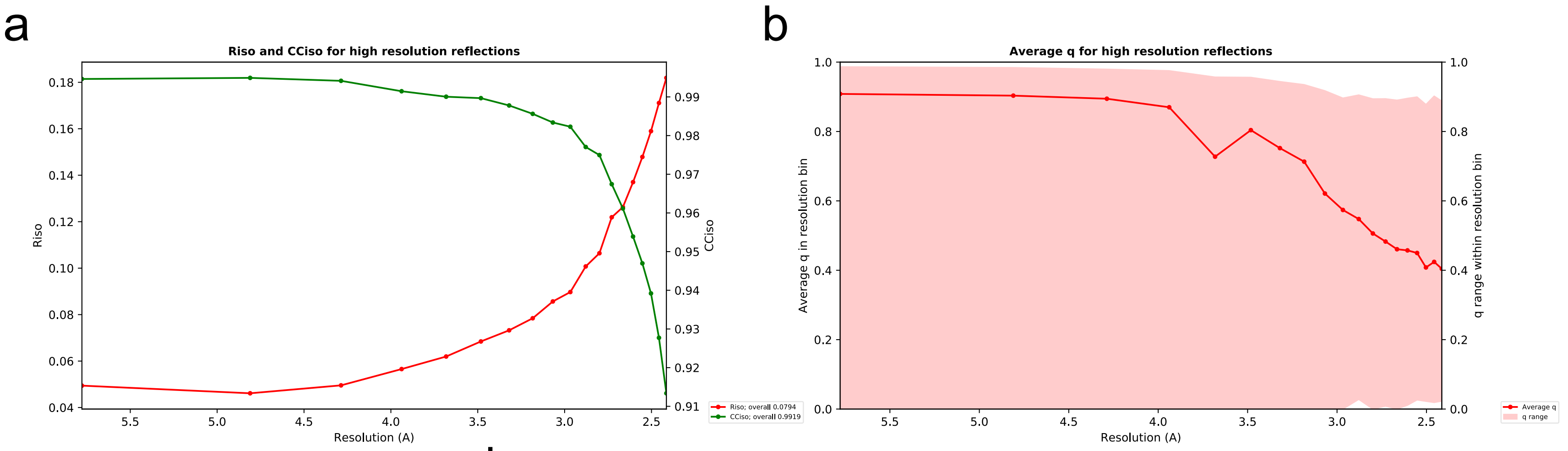


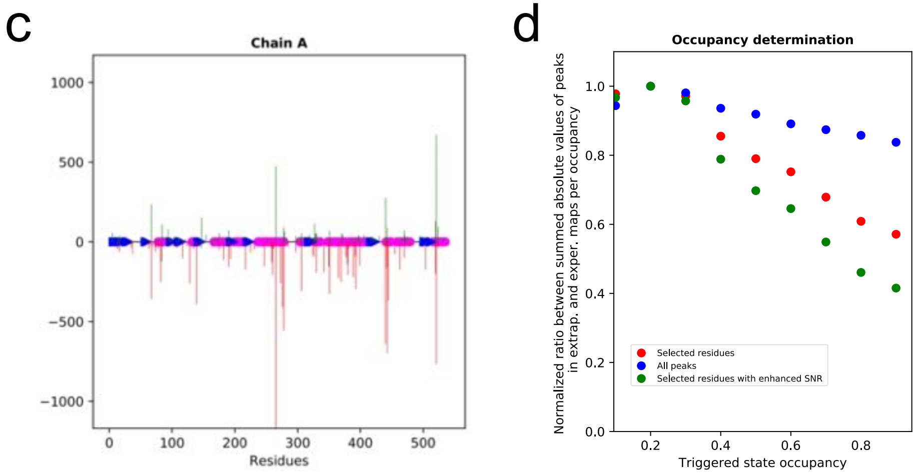


e

| Type of acidic residue | | Density visible at 0.8 σ in $2qmF_{ext}^{2vjb-2vja} - DFc$ maps | |
|---|---|---|---|
| | | Monomer A | Monomer B |
| Aspartate | Number of residues | 6 | 9 |
| | Residues number | 72, 93, 172, 182, 326, 397 | 72, 93, 172, 182, 326, 377, 380, 381, 397 |
| | Residues consistently preserved from damage | 72, 93, 172, 182, 326, 397 | |
| Glutamate | Number of residues | 7 | 8 |
| | Residues number | 5, 82, 92, 443, 462, 499 | 5, 73, 92, 139, 163, 240, 306, 443, 455, 462 |
| | Residues consistently preserved from damage | 5, 92, 443, 462 | |

**Supplementary Fig. 22 |** Data quality assessment and analysis of structural differences in the Shoot-and-Trap test case using the data sets collected at 100K. **a,** $R_{iso}$, $CC_{iso}$ values indicating the isomorphism between the 2VJA and 2VJB data sets. **b,** *q*-weight in function of the resolution. **c,** Distribution of Fourier difference peaks throughout chain A of the asymmetric unit. **d,** Occupancy determination using the difference-map method; occupancy of the triggered state is predicted to be 0.2. **e,** Analysis of X-ray induced decarboxylation of acidic residues throughout the two chains (A, B) in the asymmetric unit.

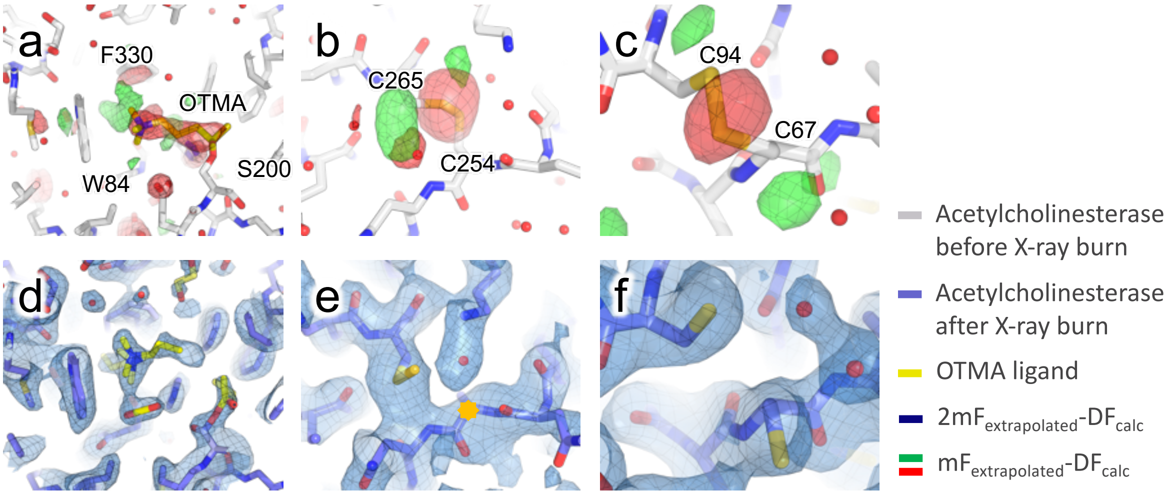


**Supplementary Fig. 23 |** Fourier difference and extrapolated electron density maps for the Shoot-and-trap experiment performed on the crystalline complex of acetylcholinesterase with a non-hydrolysable substrate analogue at 100 K. **a-c,** Specific radiation damage, manifested by negative peaks in the $q$-weighted $F_{obs}^{2VJB} - F_{obs}^{2VJA}$ Fourier difference map, is observed on the substrate analogue, carboxylate groups and disulfide bridges, respectively, indicating cleavage of the covalent bond to the protein, decarboxylation and rupture of disulfide bridges. Positive peaks are also seen that point to a reorientation of radiolysis-products in the active site (CO2, carbocholine, ethanal) and throughout the protein ($CO_2$, cysteines). **d,** the $q$-weighted $2mF_{extrapolated}$-$DF_{calc}$ extrapolated map clarifies the outcome of the radiolysis at 100 K of the covalently bounded substrate analogue, revealing how $CO_2$, carbocholine and ethanal reorient in the active site. **e-f,** Extrapolated maps also clearly evidence X-ray-induced rupture of disulfide bridges.

**Supplementary Table 1 |** Types of ESFAs calculated by Xtrapol8 and corresponding map coefficients.

| Type | Extrapolated structure factors | Map Coefficients | |
|---|---|---|---|
| | | **2F<sub></sub>** | |
| | | **$2F_{extrapolated}$-$F_{calc}$ type** | **$F_{extrapolated}$-$F_{calc}$ type** |
| **$F_{extr}$** | $F_{extr}= \alpha \times(F^{trig}_{obs}- F^{ref}_{obs})+F^{ref}_{obs}$ | $2m\|F_{extr}\|-D\|F^{ref}_{calc}\|, \varphi^{ref}_{calc}$ | $m\|F_{extr}\|-D\|F^{ref}_{calc}\|, \varphi^{ref}_{calc}$ |
| **$qF_{extr}$** | $qF_{extr}= \alpha \times\frac{q}{\langle q\rangle} \times(F^{trig}_{obs}- F^{ref}_{obs})+F^{ref}_{obs}$ | $2m\|qF_{extr}\|-D\|F^{ref}_{calc}\|, \varphi^{ref}_{calc}$ | $m\|qF_{extr}\|-D\|F^{ref}_{calc}\|, \varphi^{ref}_{calc}$ |
| **$kF_{extr}$** | $qF_{extr}= \alpha \times\frac{k}{\langle k\rangle} \times(F^{trig}_{obs}- F^{ref}_{obs})+F^{ref}_{obs}$ | $2m\|kF_{extr}\|-D\|F^{ref}_{calc}\|, \varphi^{ref}_{calc}$ | $m\|kF_{extr}\|-D\|F^{ref}_{calc}\|, \varphi^{ref}_{calc}$ |
| **$F_{genick}$** | $F_{genick}= \alpha \times(F^{trig}_{obs}- F^{ref}_{obs})+F^{ref}_{obs}$ | $m^{ref}\|F_{genick}\|, \varphi^{ref}_{calc}$ | $m^{ref}\|F_{genick}\|-D\|F^{ref}_{calc}\|, \varphi^{ref}_{calc}$ |
| **$qF_{genick}$** | $qF_{genick}= \alpha \times\frac{q}{\langle q\rangle} \times(F^{trig}_{obs}- F^{ref}_{obs})+F^{ref}_{obs}$ | $m^{ref}\|qF_{genick}\|, \varphi^{ref}_{calc}$ | $m^{ref}\|qF_{genick}\|-D\|F^{ref}_{calc}\|, \varphi^{ref}_{calc}$ |
| **$kF_{genick}$** | $qF_{genick}= \alpha \times\frac{k}{\langle k\rangle} \times(F^{trig}_{obs}- F^{ref}_{obs})+F^{ref}_{obs}$ | $m^{ref}\|kF_{genick}\|, \varphi^{ref}_{calc}$ | $m^{ref}\|kF_{genick}\|-D\|F^{ref}_{calc}\|, \varphi^{ref}_{calc}$ |
| **$F_{extr\_calc}$** | $F_{extr\_calc}= \alpha \times(F^{trig}_{obs}- F^{ref}_{obs})+F^{ref}_{calc}$ | $2m\|F_{extr\_calc}\|-D\|F^{ref}_{calc}\|, \varphi^{ref}_{calc}$ | $m\|F_{extr\_calc}\|-D\|F^{ref}_{calc}\|, \varphi^{ref}_{calc}$ |
| **$qF_{extr\_calc}$** | $qF_{extr\_calc}= \alpha \times\frac{q}{\langle q\rangle} \times(F^{trig}_{obs} - F^{ref}_{obs})+F^{ref}_{calc}$ | $2m\|qF_{extr\_calc}\|-D\|F^{ref}_{calc}\|, \varphi^{ref}_{calc}$ | $m\|qF_{extr\_calc}\|-D\|F^{ref}_{calc}\|, \varphi^{ref}_{calc}$ |
| **$kF_{extr\_calc}$** | $F_{extr\_calc}= \alpha \times\frac{k}{\langle k\rangle} \times(F^{trig}_{obs}- F^{ref}_{obs})+F^{ref}_{calc}$ | $2m\|kF_{extr\_calc}\|-D\|F^{ref}_{calc}\|, \varphi^{ref}_{calc}$ | $m\|kF_{extr\_calc}\|-D\|F^{ref}_{calc}\|, \varphi^{ref}_{calc}$ |

$\alpha$=1/occupancy; $F^{trig}_{obs}$, $F^{ref}_{obs}$: observed structure factors amplitudes associated to triggered and reference state, respectively; $F^{ref}_{calc}$: calculated structure factors associated to reference state; $m^{ref}$: figure of merit extracted from the reference data.
